# Insights into the spool-like architecture and infection strategy of the archaeal virus SEV1

**DOI:** 10.1101/2024.12.18.629138

**Authors:** Haonan Zhang, Haina Wang, Yan Li, Yunxuan Fan, Zhenfeng Zhang, Hongyu Chen, Kai Song, Li Huang, Ping Zhu

## Abstract

Archaeal viruses are well known for their diverse morphologies and extreme stability. In this study, we used cryo-electron tomography to analyze the structure of SEV1 and its infection strategies in its native state. The results show that SEV1 nucleocapsid adopts a ‘coil-stacking’ architecture which displays a degree of flexibility. VP4, whose homologues are widespread in the thermo-acidic environment globally, is identified as the major capsid protein and binds genomic DNA forming a “beads-on-a-string” arrangement. Simulations in various extreme environments indicate that the envelope of SEV1 is crucial to the thermostability. SEV1 infects the host by membrane fusion revealed by the membrane fusion assay. The infected cell undergoes cytoplasm condensation to form a “viral factory”, leading to the successive production of nascent virions. A series of assembly intermediates of SEV1 are identified revealing an integrated picture of the virus assembly process. The nascent virions are found to be released through virus-associated pyramids (VAPs), composed of unique proteins encoded by SEV1 distinct from other known VAP proteins. Our study provides novel insights to the survival strategies of SEV1, a flexible and enveloped archaeal virus, by using the unique “coil-stacking” architecture and the characteristic infection strategies.

## Introduction

Archaeal viruses are frequently found in extreme habitats, including hot springs, hydrothermal vents, and salt lakes, and often exhibit an unusual morphology, such as a filament, a spindle, a bottle, a droplet, a sphere or a pleomorphic shape [1]. In the past decade, cryo-electron microscopy (cryo-EM) has been increasingly employed to reveal the architectural details of archaeal viruses, shedding light on their ability to survive under harsh conditions [2-5]. Filamentous and rod-shaped archaeal viruses, which are typically classified into four viral families, i.e., Rudiviridae, Lipothrixviridae, Tristromaviridae and Clavaviridae [6], are most studied, likely due to their structural rigidity and high degree of symmetry. As revealed by the near-atom resolution structure, the naked virion of *Sulfolobus islandicus* rodshaped virus 2 (SIRV2), a member of Rudiviridae, contains a linear genomic DNA tightly wrapped and transformed into A-form by the major capsid protein (MCP) [7]. As a result of the extensive hydrophobic interactions within and among the MCP dimers, the encapsidated viral genomic DNA is entirely separated from solvent [7]. By contrast, the *Acidianus* filamentous virus 1 (AFV1), an enveloped lipothrixvirus, lacks the hydrophobic protein-protein interaction across the helical turns of the virion, and the genomic DNA is protected by the lipid envelope [8]. Exploration of the structures of archaeal viruses has led to the notion that packing of A-form DNA is a common strategy of archaeal viruses to stabilize DNA in extreme environments, as it has been observed in *Pyrobaculum* filamentous virus 2 (PFV2) [6] and icosahedral *Sulfolobus* polyhedral virus 1 (SPV1) [3], in addition to other rudiviruses [4, 7] and lipothrixviruses [4, 8, 9]. Filamentous *Aeropyrum pernix* bacilliform virus 1 (APBV1) packs its double-stranded DNA (dsDNA) as a left-handed superhelix, and its helical capsid tightly packed through hydrophobic interaction likely contributes to the thermostability of the virion [10]. Hydrophobic protein-protein interaction in the capsid is presumably also responsible for the thermostability of *Sulfolobus* spindle-shaped virus 19 (SSV19) [5]. However, how genome is packaged in the structurally flexible archaeal viruses remains largely unknown.

Archaeal viruses are also known to have evolved various infection strategies. SIRV2 initiates infection by binding to the pilus-like filament on the host with its terminal appendages [11], while the enveloped halophilic archaeal virus Haloarchaeal pleomorphic virus 6 (HRPV-6) infects its host cell by membrane fusion [12]. Membrane fusion is presumably also employed by spindle-shaped virions of *Sulfolobus* monocaudavirus 1 (SMV1) for host entry [13]. Distinct strategies appear to be used by archaeal viruses during the process of virions assembly and egress. Virions of the spindle-shaped virus SSV1 are assembled underneath the host cell membrane, and exit the host while acquiring the lipid membrane from the host through budding [14]. In contrast, despite their morphological differences, filamentous SIRV2 [15], *Sulfolobus islandicus* filamentous virus (SIFV) [16] and icosahedral *Sulfolobus* turreted icosahedral virus (STIV) [17] are assembled into mature virions inside the host cell and released from the host cell through the polygonal virus-associated pyramids (VAPs), a special structure formed on the host membrane. However, the knowledge of the life cycles of enveloped and structurally flexible archaeal viruses is still fragmentary.

*Sulfolobus* ellipsoid virus 1 (SEV1), an archaeal virus isolated from an extremely acidic hot spring in Costa Rica [18], is the sole member of the archaeal virus family *Ovaliviridae* [19] and a morphologically flexible virus enveloped by a lipid membrane [18]. The nucleocapsid of SEV1 exhibits a morphology characterized by 16 regularly spaced striations, suggesting a multi-layered disc-like architecture likely formed by intricate winding of a linear viral nucleoprotein filament [18]. Interestingly, as an enveloped virus, the SEV1 virion acquires its lipid envelope during maturation in the host cytoplasm and is released from the host cell through a VAP [18]. Nevertheless, the detailed architecture and infection process of the SEV1 virion, as well as their contribution to the ability of the virus to thrive in extreme environments remain elusive.

In this study, we characterized the structure of SEV1 and explored its infection process by using a combination of cryoelectron tomography (cryo-ET), cryo-focused ion beam (cryo-FIB), tomogram segmentation, sub-tomogram averaging (STA), artificial intelligence (AI)-based structural prediction, and biochemical analysis. By cryo-ET 3-D reconstruction, we revealed a remarkable ‘coil-stacking’ mode for the packaging of the SEV1 genomic DNA in association with the major capsid protein (MCP)VP4, which presumably is responsible for the general flexibility and local regularity of the virus particle. By using cryo-FIB sectioning and cryo-electron tomography in combination, we obtained an integrated picture of the entire infection process of SEV1 in the native state. We found that SEV1 enters the host cell by destructing S-layer and membrane fusion. The infected host cells undergo cytoplasm condensation to form a “viral factory”, and the mature virions are released through VAPs constructed of a novel VAP protein. Our results provide significant insights into the architectural principle and infection process of an enveloped and structurally flexible archaeal virus and its adaptation to living in the extreme habitat.

## Results

### The coil-stacking architecture of the SEV1 nucleocapsid

SEV1 virions were cultured, purified and subjected to cryo-ET 3-D structure determination. The purified SEV1 virions are ellipsoidal and ovoid, measuring around 101.2 ± 5.5 nm in length and 79.7 ± 4.4 nm in width, and show no symmetric features (Fig. 1 A and B, Fig. S1A). AI-assisted segmentation on the reconstructed tomogram reveals that an intact SEV1 virion consists of three major parts, i.e., a nucleoprotein core, a ~5.1-nm-thick membrane envelope, and ~7.3-nm-long protruding spikes (Fig. S1A). Spikes are densely located on the envelope, which encloses the nucleoprotein core of a multi-layered architecture (Fig. S1B and C). As we previously observed [18], the virions of SEV1 display a distinct pattern of striation in the side view and coils in the top view (Fig. 1A), although irregularity in nucleoprotein stacking was observed both in purified virions and those within the infected cells (Fig. 2A-B, Fig. S2). The nucleocapsid core in some virions appears to be broken into two parts, mostly at the middle, with one part being flipped (Fig. 2A-C). Virions with an irregular shape were also observed, and in some of them the coil-stacking pattern was maintained in separate parts of the nucleocapsid core (Fig. 2C-D).

**Figure 1:**
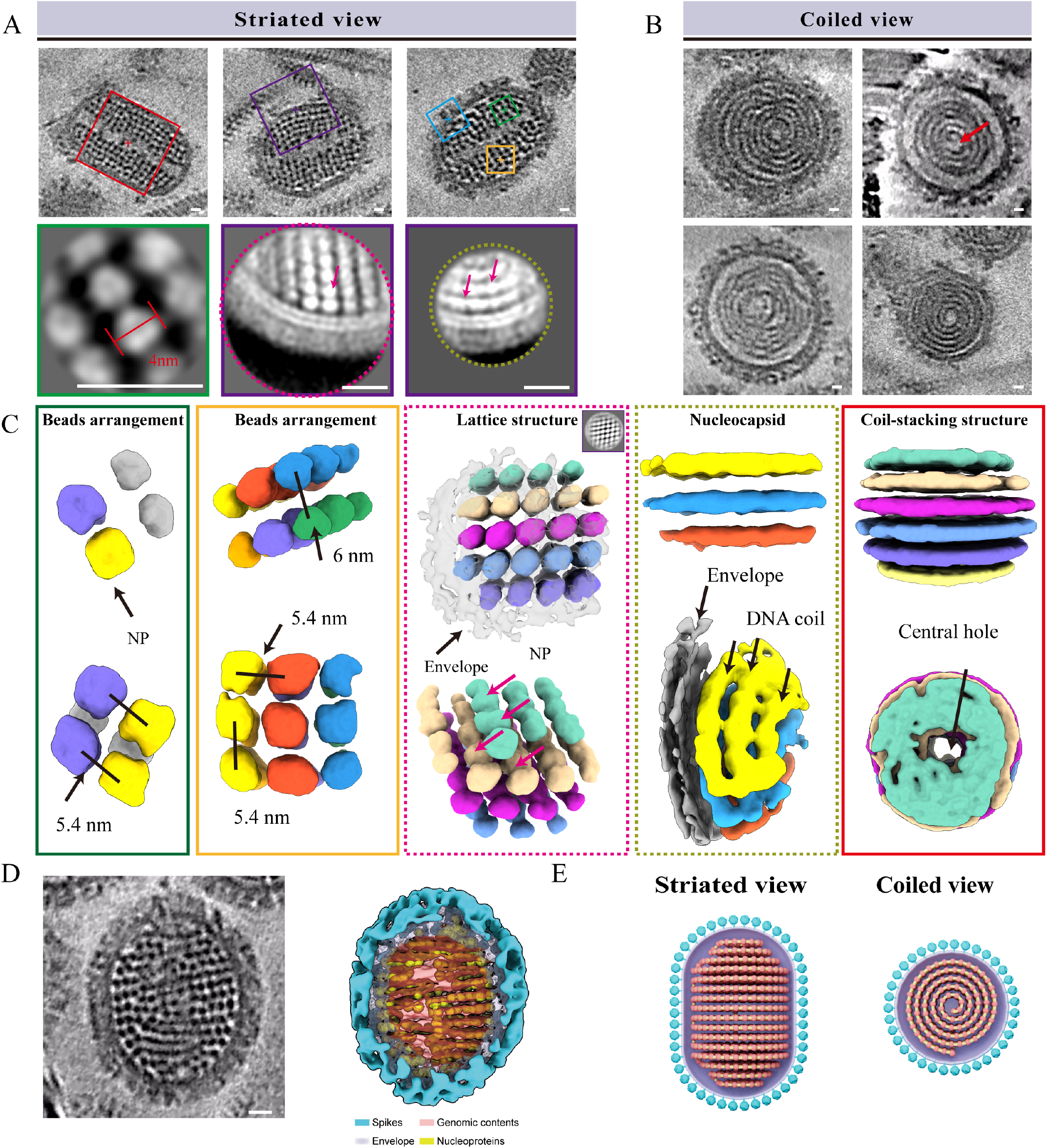
Structural characterization of SEV1 by cryo-ET. **(A)** Top: the striated view of tomogram slices of three representative SEV1 virions showing the box selection strategy. The boxes in different sizes and colours show variant strategies of sub-tomogram averaging by placing the center of sub-volumes at different locations, showing as follows: green, centering between two beads in the same disc layer; yellow, centering between two regular disc layers; red, centering in the central hole; and purple, centering beneath the envelope spike. Bottom: the slices of the averaged maps using different boxing (green and purple) and masking strategies. The red arrows in the purple box highlight the ‘bead’ (middle) and ‘strings’ (right), respectively. **(B)** The coiled view slices of representative SEV1 virions. The central hole of the SEV1 virion is indicated by a red arrow. **(C)** Structures generated from the sub-tomogram averaging (STA) using the box selection strategy as displayed in (A) and viewed from two angles (top and bottom). The colour scheme corresponds to the boxes in (A). **(D)** Left: a tomographic slice of the reconstructed SEV1 virion. Right: NPs mapped back to the SEV1 virion. The spikes, envelope and the genomic contents are the segmented structures, while the NPs are the averaged map that remapped to the segmented genomic contents. The beads well-merged with the segmented genomic contents are highlighted in brown. **(E)** The model of SEV1 virion displayed by striated view and coiled view. The nucleoprotein filament in the ‘beads-on-a-string’ pattern is composed of NPs (red) densely distributed on the DNA (yellow). The nucleoprotein filament coils and stacks to form the nucleocapsid which is enveloped by the lipid membrane (purple). The blue balls anchored in the lipid membrane represent the spikes of SEV1. Scale bar: 10 nm.

**Figure 2:**
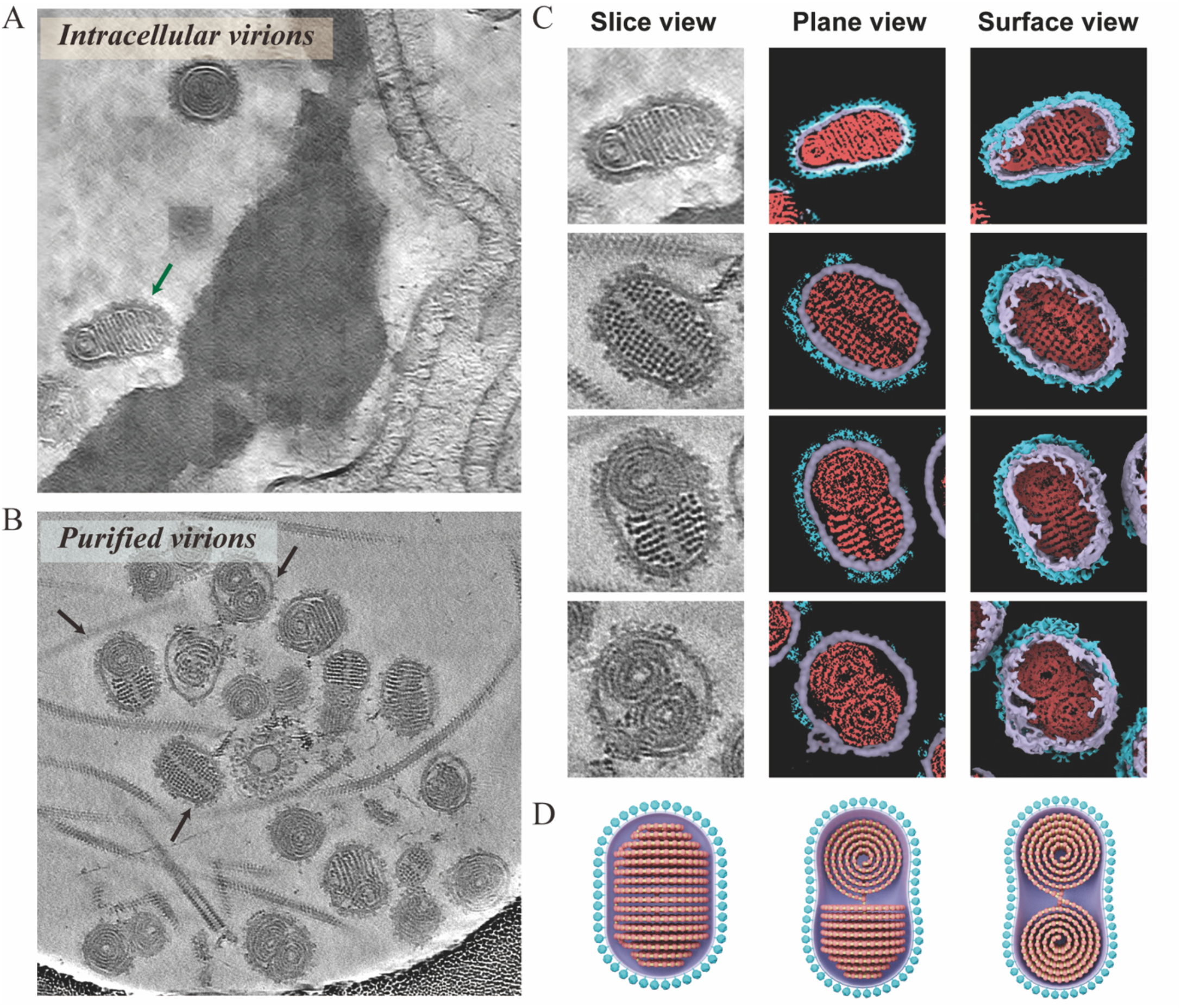
Variations and flexibility of the SEV1 nucleocapsid. **(A)** A tomogram slice showing SEV1 virions assembled inside the host cell of *Sulfolobus* sp. A20. A SEV1 virion with partially rotated nucleocapsid is indicated by a green arrow. **(B)** A representative slice of purified SEV1 virions showing variations in nucleocapsid. Three virus particles with unusual nucleocapsid structures are indicated by black arrows. **(C)** The slice, plane and surface views of the SEV1 virions identified in (A) and (B). **(D)** Models of the three typical nucleocapsid structures.

Notably, plentiful ‘beads’ are densely arranged in a highly regular pattern in the nucleocapsid, as shown in the tomograms (Fig. 1A), suggesting the formation of nucleoprotein (NP) complexes with the appearance of ‘beads on a string’. To look further into these regularly distributed ‘beads’, i.e., NPs, we performed sub-tomogram averaging (STA) on boxed sub-volumes containing a number of beads (Fig. 1A). Firstly, we averaged the sub-tomogram volumes in a box size of ~16×16×16 nm by putting the box center between two beads on the same layer (Fig. 1A, top panel, green box). Around 500 sub-volumes were manually picked and averaged. In the average map, six beads could be clearly observed (Fig. 1A, bottom panel, green box). The inter-coil and the lateral distances between two adjacent NPs are both measured 5.4 nm, suggesting a highly regular distribution of the NPs (Fig. 1C, green box). We then expanded our attempt to average the sub-tomogram volumes with a larger box size (~19×19×19 nm) but putting the box center between two adjacent disc layers (Fig. 1A, yellow box). The average map shows a two-layer arrangement of NPs. Each layer contains 9 regularly distributed NPs (Fig. 1C, yellow box). The distance between neighboring NPs on the same layer is again measured 5.4 nm, identical to that observed above in the analysis of six beads, while the distance between two adjacent layers is ~6 nm (Fig. 1C, yellow box).

It is challenging to obtain the arrangement pattern of NPs in the box containing more NPs due to the high flexibility of viral particles. Unexpectedly, however, we partially archived this goal when we attempted to acquire the structure of the protruding spike by STA. In this effort, we defined a normal vector perpendicular to the envelope of the virus and sampled (boxed out) the spikes evenly. The given initial orientation of individual spike facilitated a better alignment of the boxed NPs-containing sub-volume boxes centering on the nucleocapsid beneath the corresponding envelope spike (Fig. 1A, purple box). We found that different features of the nucleocapsid could be revealed when we averaged the sub-volumes with differently-sized masks. When a large mask (Fig. 1A, dashed magenta circle) was used, the averaged map displayed a lattice-like arrangement of beads in the capsid, especially right beneath the envelope (Fig. 1C, dashed magenta box). Intriguingly, when a smaller mask (Fig. 1A, dashed brown circle) was used, discrete strands of nucleoprotein filaments became apparent on each disc layer, forming a ‘DNA coil’ (Fig. 1C, dashed brown box). These findings are consistent with the features observed in tomographic slices (Fig. 1A, purple boxes).

As shown in Fig. 1A (red box), the coiled structure appears to contain a central channel, as proposed previously [18]. We then performed STA by putting the box center on the innermost coil as indicated by the red arrow in Fig. 1B, which corresponds to the area in red box in Fig. 1A. As expected, the averaged map revealed a pronounced stacking of discs with a conspicuous channel, perpendicular to the discs, at the center of the coiled structure and spanning the central region of the entire nucleocapsid (Fig. 1C, red box).

When we mapped all of the NP beads used in the sub-volume averaging (averaged in green box) to the virion (Fig. 1D, left), it becomes clear that the SEV1 nucleocapsid comprises stacked discs, each of which contains nucleoprotein coils with the ‘beads-on-a-string’ appearance, and is enclosed in a spike-decorated envelope to form a complete virion (Fig. 1D, right).

To determine how the linear genomic DNA threads between the discs, we looked closely into points where two adjacent discs were joined in the tomographic slices. We found that the contact points are close to either the central channel or the outer edge of the nucleocapsid, suggesting that the nucleoprotein filament switches the direction of coiling between layers. In other words, if the filament coils inward to the central channel in one disc, it would coil outward in the adjacent disc (Fig. S3, white arrow). It appears that this coiling strategy permits tight stacking of the discs and genome packaging in the SEV1 nucleocapsid.

Surprisingly, the SEV1 virions display pronounced variation in nucleocapsid structure. Virions containing a partially flipped nucleocapsid, referred to as special virions hereafter, were observed in both the infected *Sulfolobus* sp. A20 cell and the purified virion preparations (Fig. 2). The presence of a virion with a nucleocapsid showing both striations and coils in the infected cell excludes the possibility of the deformation of the virus particles during virus preparation (Fig. 2A). Interestingly, the two parts of nucleocapsid in these virions are related by ~90° rotation between them or are aligned side by side facing the same direction (Fig. 2C). In addition to these two typical structural variations, other anomalies in nucleocapsid were also observed, suggesting the structural flexibility of SEV1 (Fig. S3). Despite its flexibility, the irregular SEV1 nucleocapsids still exhibit the characteristic striation and coiling, and thus maintains the coil-stacking architecture.

### VP4, the major capsid protein of SEV1

SEV1 encodes four structural proteins [8]. To identify the protein(s) in the nucleocapsid, we removed the envelope from the virus by treatment with 1% NP-40 (Fig. 3A-B). The resulting envelope-free virus fraction contained mainly a single protein of ~12 kDa in size (Fig. 3A), which was identified as VP4 by mass spectrometry (Table S1). Therefore, we conclude that VP4 is the protein component of the observed beads (Fig. 1A) in the nucleoprotein filament and the MCP of SEV1. We then subjected the virus particles to multiple cycles of freeze and thaw to break the envelope. The nucleocapsids released from the partially disassembled virions were examined by cryo-ET (Fig. S4A). The SEV1 nucleoprotein filament was found significantly and uniformly thicker than naked DNA, leaving out hardly any exposed DNA, suggesting that the genomic DNA is entirely coated with VP4 (Fig. S4A). In agreement with this observation, the SEV1 nucleocapsid was highly resistant to cleavage by micrococcal nuclease (MNase), as expected from the lack of access of the enzyme to the virus DNA as the result of tight binding by VP4 (Fig. S4B).

**Figure 3:**
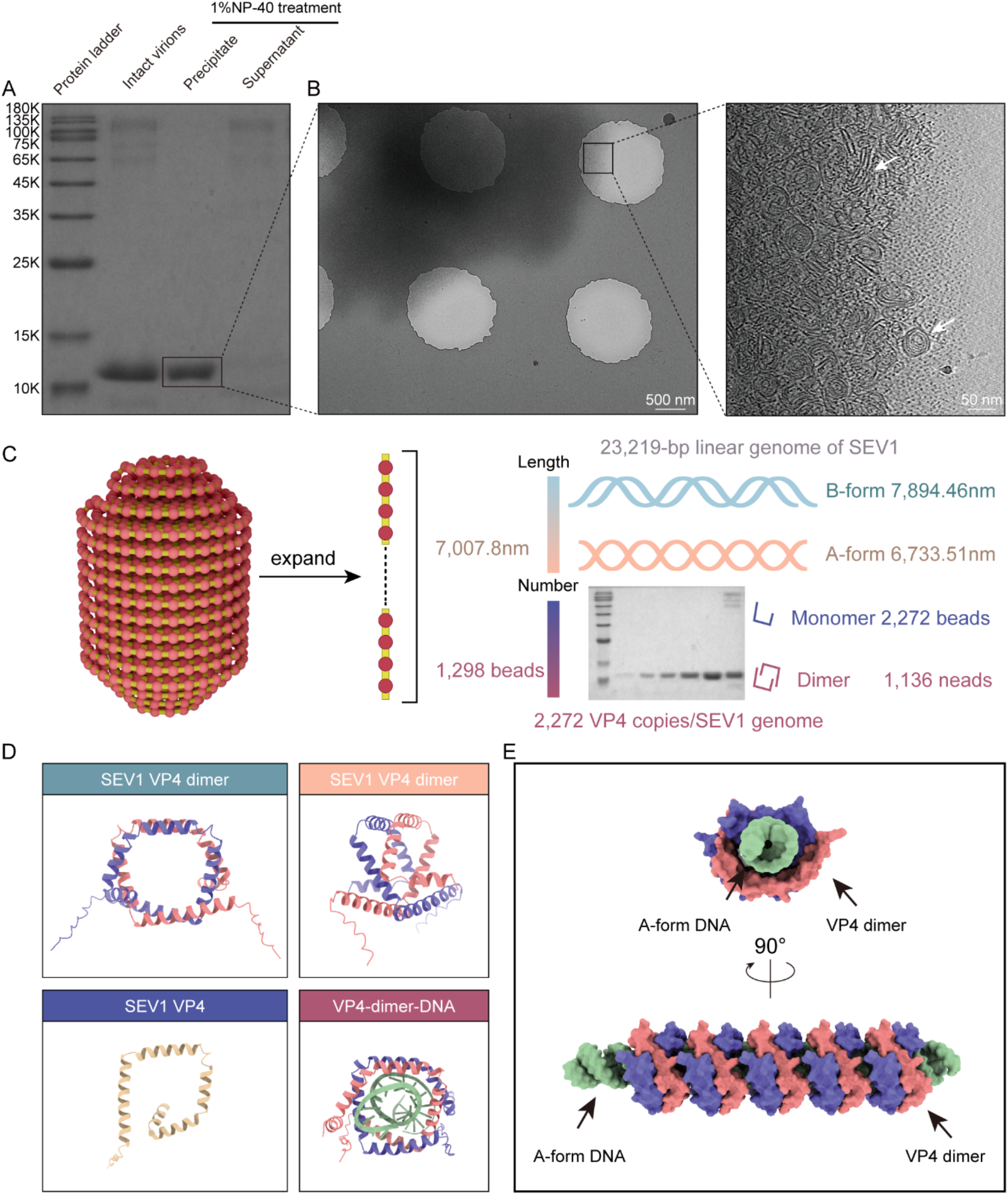
Identification of the nucleocapsid protein VP4 and the interaction of VP4 with the virus DNA. **(A)** of the treated SEV1 virions. **(B)** Cryo-EM of the NP-40-treated SEV1 virions. Low magnification image shows large aggregates covering the hole of the grid square. Zoom-in tomographic slice of the large aggregate displays nucleocapsids in coil-like and disc-stacking views. **(C)** Schematic diagram of A-form DNA wrapped by VP4 dimers. Based on the proposed model, the SEV1 genomic DNA is in the A-form and covered by VP4 dimers. **(D)** Structural prediction of VP4 and the VP4-DNA complex by AlphaFold. **(E)** Model of SEV1 DNA packaging by VP4 dimer, as displayed in top view and side view.

Further, we performed a semi-quantitative analysis of the amount of VP4 in SEV1 virions by SDS-PAGE and subsequent Coomassie blue gel staining using the envelope-free virus fraction, whose protein concentration was predetermined, as a reference (Fig. 3C). The amount of the genomic DNA in the same virus preparation as that used for the VP4 quantification was also determined. Based on these measurements, we derived a stoichiometry of 2,272 VP4 molecules per virus genome (Fig. 3C). On the other hand, the number of NP beads in a SEV1 virion was estimated to be 1,298 based on the structural model of the virus. So, the VP4/bead ratio is 1.75 or close to 2, suggesting that VP4 binds the genomic DNA as a dimer. SEV1 possesses a linear genome of 23,219 bp, which corresponds to a B-form DNA of ~7,894 nm or A-form DNA of ~6,734 nm in length. According to our structural model, the dsDNA in one SEV1 virion is estimated to be 7,008 nm long (Fig. 3C), and likely adopts the A-form, as found in SPV1, which also harbors a coiled nucleocapsid [3], and several other archaeal viruses [4, 7, 8].

To gain clues to the interaction between VP4 and the virus genomic DNA, we predicted the structure of the VP4 dimer and the VP4-DNA complex using AlphaFold2 and AlphaFold3 (Fig. 3D). The predicted structure shows that each monomer in the VP4 dimer adopts a structure containing 5 helices which form a central channel. Two VP4 monomers stack to each other to form a VP4 dimer containing a central hole, through which the DNA passes (Fig. 3D). The predicted VP4-DNA complex structure is consistent with the model that the genomic DNA in the A form threads through VP4 dimers, leading to the formation of the SEV1 nucleocapsid (Fig. 3E). This model agrees well with that proposed for the SPV1 nucleocapsid [3], suggesting that the coil-stacking conformation is conserved among archaeal viruses, regardless of their morphology, as a strategy to stabilize the virus genome in harsh habitats.

A psi-blast search with the amino acid sequence of SEV1-VP4 in the non-redundant database of GenBank resulted in only a single homologous sequence, which is from the metagenome-assembled genome of *Sulfolobus* Bepepu virus (SBV) from a Japanese acidic hot spring [20]. We then attempted a search against IMG/VR v4.1, an expanded database of uncultivated virus genomes within a framework of extensive metadata [21], and retrieved 50 sequences with associated habitat pH and temperature data (3 iterations, e-value <10^−5^) (Table S2). A majority (48) of the identified homologues are from acidic hot springs and sediments in YNP, and the remaining two sequences are from an acidic hot spring in Japan and a deep-sea sediment sample in California Gulf (Mexico). SEV1-VP4 homologues comprise three clades in the phylogenetic tree, suggesting evolutionary divergence of these putative MCPs and the viruses that possibly encoding them (Fig. S5). Further, we predicted the structures of 10 VP4 homologues representing the three clades (Fig. S5A), and found that they all share a well conserved structure (Fig. S5C). Therefore, if these VP4 homologues are the MCPs of viruses, these viruses may employ the same genome packaging mechanism as that of SEV1.

Among the identified SEV1-VP4 homologues, only GP-06, encoded by SBV, is from the enrichment culture containing virus-like particles resembling SEV1 in morphology [20], and the remaining homologues are derived from the viral metagenomic assembled contigs without clear classification. Sequence alignment shows that SEV1-VP4 and SBV-GP-06 share a similar alpha-helix arrangement (Fig. S6). By AlphaFold prediction, SEV1-VP4 and SBV-GP-06 are structurally similar and are able to form the central hole via 5 helices (Fig. S5C), as proposed in the ‘beads-on-a-string’ model for the SEV1 nucleocapsid (Fig. 3C). Surprisingly, SEV1-VP4 shares no sequence similarity with SPV1-VP1, as the psi-blast with the SEV1-VP4 sequence failed to retrieve SPV-VP1 in the GenBank, although SEV1 and SPV1 have similar coil-stacking nucleocapsids. Despite their lack of similarity at the amino acid sequence level, SEV1-VP4 resembles SPV1-VP1 in secondary structure as both proteins consist of five alpha-helices [3] and package DNA in the ‘beads-on-a-string’ pattern.

### Structural stability of SEV1 to pH and temperature

To learn more about the structural stability of SEV1, we exposed the virus to different pHs and temperatures, and performed cryo-ET analysis on the treated virus particles (Fig. 4A). We first incubated the purified SEV1 virions at various pHs at 23°C. As shown in Fig. 4A, the nucleocapsid of SEV1 exhibits a normal coil-stacking architecture at pH 3, the optimal pH for the growth of the host, and remains largely unchanged at pH 5-7. However, occasional breaks of the envelope, accompanied by the slight leakage of the nucleoprotein filament, are detectable at pH7. The lipid envelope starts to break down at pH 9, and is severely damaged at pH 11 and 13, leading to the disassembly of the NPs and the fragmentation of the viral DNA. On the other hand, the nucleocapsid becomes more condensed at pH 1 (Fig. 4A) than at pH 3. Interestingly, the SEV1 virion, especially the viral spikes, appear to undergo significant morphological changes in response to changes in pH at the very low end (Fig. 4C). For example, the length of the spike extends from 7 nm to 17 nm as pH decreases from 1 to 0.5. We then incubated the SEV1 virions at different temperatures at a fixed pH of 3. We found that SEV1 is highly resistant to high temperatures, and is able to retain an intact nucleocapsid structure at 100°C (Fig. 4A, bottom right panel and Fig. S7). These results suggest that SEV1 virions are extremely stable at acidic pH (pH 1-5) and high temperature (up to 100°C), but are not as stable under alkaline conditions.

**Figure 4:**
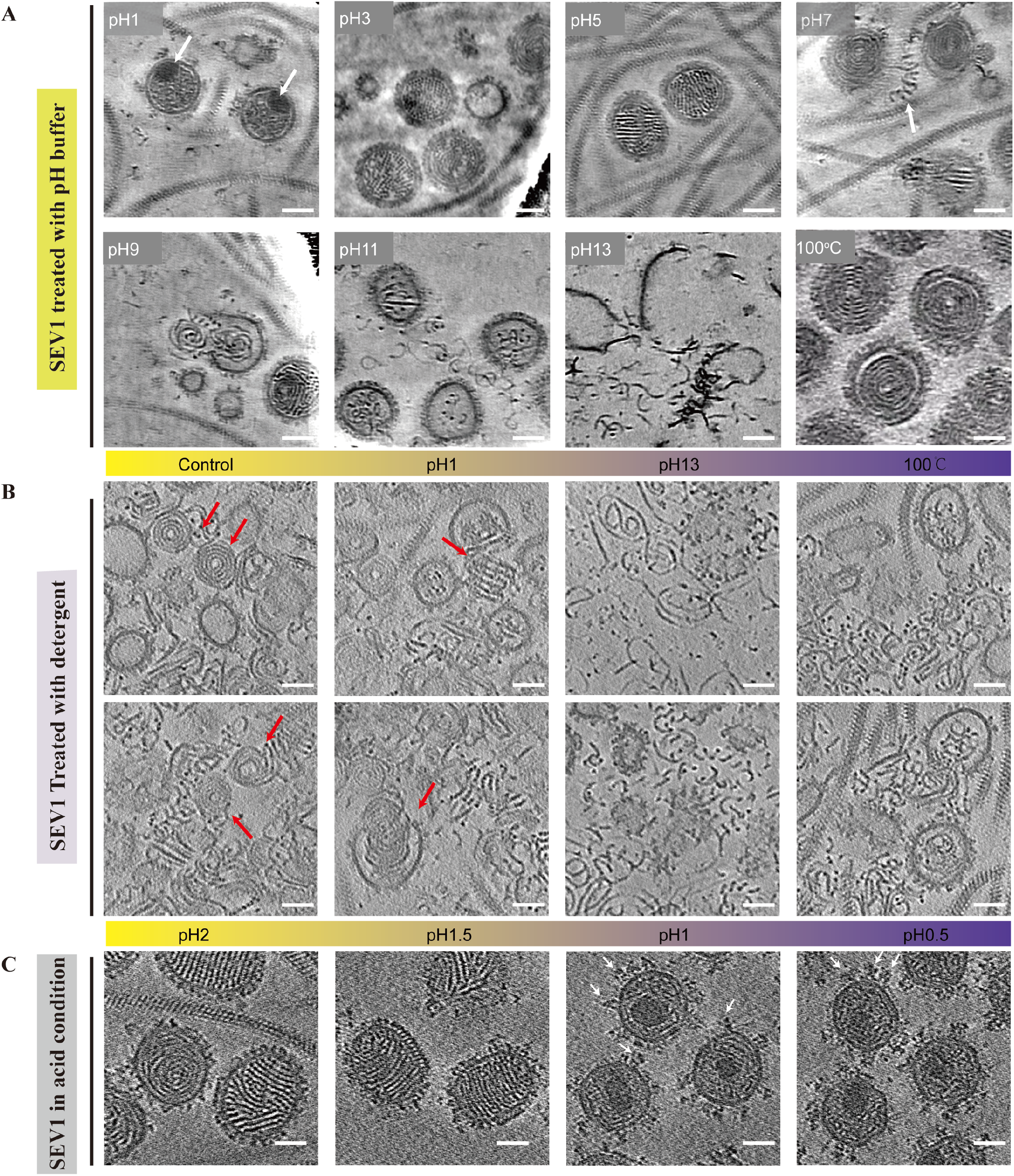
Effect of pH and temperature on the structural stability of SEV1. **(A)** SEV1 virions incubated at various pHs (room temperature) or at 100°C (pH3). **(B)** Triton-treated SEV1 virions incubated at various pHs or at 100°C. **(C)** SEV1 virions incubated at highly acid conditions (pH 0.5-2).

To explore the contribution of the lipid envelope to the stability of SEV1, we treated the virus particles with 0.1% Triton X-100 to obtain the envelope-less nucleocapsid of SEV1, and performed the same stability analysis on the naked SEV1 nucleocapsid as that on the intact virions (Fig. 4B). The naked nucleocapsid is stable, displaying the striated and coil-like pattern in the tomogram slice at pH 3. However, unlike the enveloped virions, the naked virions do not condense and maintain a normal coil-stacking architecture at pH 1, suggesting that the envelope may serve a role in nucleocapsid condensation observed at the extremely low pH. On the other hand, the naked nucleocapsid remains stable at pH 5, disassembles to a gradually increasing extent when pH is raised from 5 to 9, and collapses completely at higher pHs (Fig. 4B). Therefore, the pH stability of the naked nucleocapsid is similar to that of the enveloped virion. Unlike the enveloped SEV1 virion, which is stable to temperature up to at least 100°C, the naked nucleocapsid is more sensitive to temperature within the tested range from 45 to 100°C (Fig. S8). The structure of the nucleocapsid becomes increasingly loose with an increase in temperature. Notably, the naked SEV1 nucleocapsid becomes completely disordered at 100°C. It appears that the lipid envelope contributes significantly to the structural stability of SEV1 at high temperature.

### SEV1 entry into the host cell

To examine the process of virus entry into host cell by SEV1 in a native state, *Sulfolobus* cells infected with the virus were grown at 75°C and then plunging frozen using a high-temperature sample preparation method [22]. As shown in Fig. 5A, the S-layer (highlighted by a red arrow) and the cell membrane (marked by a light-blue arrow) are identifiable at the periphery of the infected cell. In addition, chain-like densities (marked by a yellow arrow) adhering to the outer side of the S-layer are observed. These densities most likely contain glycan chains since it is known that the *Sulfolobus* S-layer is heavily glycosylated [23, 24]. The glycan chain density is also clearly visible on the reconstructed tomograms of *Sulfolobus* sp. A20 cells thinned by cryo-FIB (Fig. S9). The SEV1 virions are observed attaching to the glycan chains on the exterior of the S-layer during the early stage of entry (Fig. 5A). By NetGo prediction [25], among the structural proteins of SEV1, VP1, VP2 and VP3, but not VP4, carry putative hydrolase activity, which may be involved in host recognition and the initiation of host entry by the virus (Table S3) [5]. Interestingly, a distinct gap (indicated by the red arrow) exists in the S-layer density of the infected *Sulfolobus* cell (Fig. 5A), which is continuous in the uninfected cells. Therefore, the virus attachment to the glycan chains of the host cell appears to lead to the disruption of the S-layer, permitting the contact of the virus with the cell membrane.

**Figure 5:**
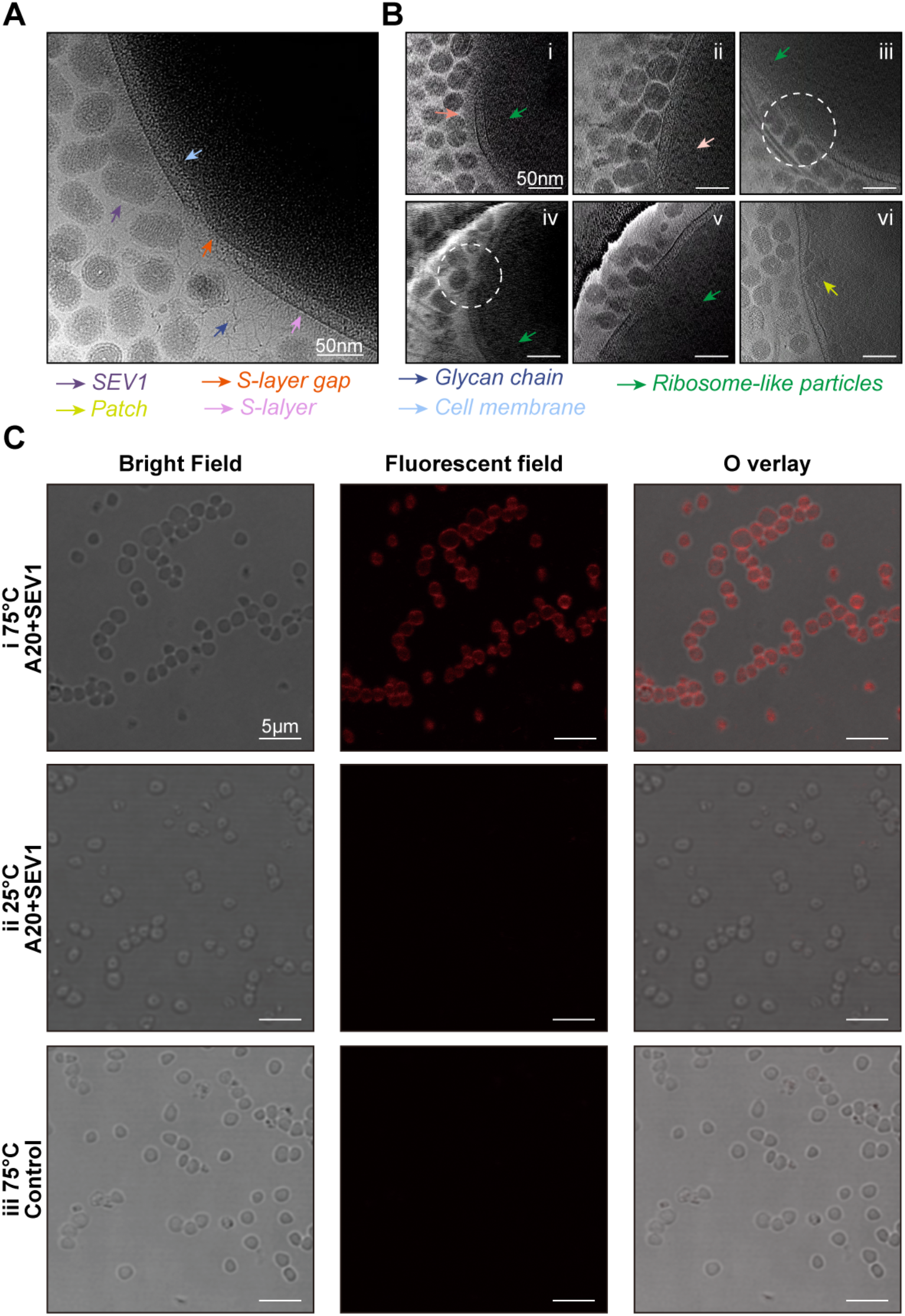
Entry of SEV1 into *Sulfolobus* sp. A20 cells. **(A)** A cryo-EM picture showing the interaction of SEV1 and the host cell envelope. The cell envelope includes S-layer, cell membrane, and cellular appendages extruding from the S-layer decorated with glycan chains. The densely hair-like glycan chains are arranged in an orderly manner and long glycan chains arranged randomly. Scale bar, 50 nm. **(B)** Sequence of events during the entry of the SEV1 virions. i. Virus particles attach to the glycan chain outside the S-layer. ii. Virus particles contact the S-layer. iii and iv. Following the interaction with the virus particles, the S-layer starts to disintegrate. The significant disruption of the S-layer is highlighted by circles. v. Virus particles penetrate the S-layer and reach the cytoplasmic membrane. vi. Patch-like structures (green arrow) are formed in the cytoplasm after the virus entry. Scale bar, 50 nm. **(C)** Virus-cell fusion assay. Fluorescent microscopy of *Sulfolobus* sp. A20 cells infected with R18 labelled SEV1 virions at 75 °C (i) and 25 °C (ii), as well as *Sulfolobus* sp. A20 cells incubated in Zillig’s basal salt at 75 °C (iii). Scale bar, 5 μm.

We then obtained snapshots of the virus-host interaction by using cryo-ET and revealed a sequence of events during virus entry. The virus first contacted the chain-like densities at the exterior of the cell envelope (Fig. 5B.i), and then penetrated the dense layer of glycan chains to get in touch with the S-layer (Fig. 5B.ii). The interaction of the virus with the S-layer resulted in pronounced structural disruption spanning the entire S-layer, allowing the virus to interact with the cell membrane (Fig. 5B.iii, Fig. 5B.iv). This is followed by the internalization of the virus (Fig. 5B.v) and the formation of patch-like structures, which presumably consist of nucleoprotein complexes [14], within the cell (Fig. 5B.vi).

To explore if the SEV1 entry involved fusion between the virus envelope and the cell membrane, we employed a virus-cell fusion assay including the use of the lipophilic fluorescent dye octadecyl rhodamine B chloride (R18) as a tracer [12]. R18-labeled virions were incubated with unlabeled *Sulfolobus* sp. A20 cells. After 1.5 hr at 75°C, the cytoplasmic membrane of the cells was completely fluorescent (Fig. 5C), indicating that R18 had diffused from the viral envelope to the cell membrane. As controls, the host cells incubated with either R18-labeled virions at 25°C, a temperature at which SEV1 was unable to infect the host cell, or in Zillig’s basal salt at 75°C exhibited no fluorescence, demonstrating that membrane fusion between SEV1 and the host cells could not occur in the absence of the virus infection. Therefore, we conclude that host entry by SEV1 involves membrane fusion.

### Virus assembly and release from the host cell

To shed light on SEV1 assembly and egress in the native state, we next investigated the virus-infected culture by cryo-FIB and cryo-ET (Fig S10A, B). The reconstructed tomograms were analyzed using AI-based deep learning technology to annotate automatically cell envelope, cytoplasmic contents, and virion components [26]. In the uninfected cell, the cytoplasmic contents are evenly distributed, and the S-layer proteins attach tightly to the cell membrane, representing a typical architecture of the *Sulfolobus* cell envelope (Fig. 6A.i). Following infection, progeny virus particles are assembled and, notably, enveloped into mature virions in the cytoplasm, where ribosomes-like particles are visible (Fig. 6A. ii). Moreover, the cryo-FIB milling and the readily visible cryo-electron tomogram with high resolution allow the virus components (i.e., spikes, envelope, and nucleocapsid) to be clearly segmented in the cellular environment (Fig. 6A, bottom panels). As described above, intracellular virions exhibiting the characteristic structural pattern of coil stacking are clearly observed (red and blue arrows in Fig. 7Aii). The purified virions and intracellular virions are measured 101.2 ± 5.5 nm long and 79.7 ± 4.4 nm wide and 99.6 ± 6.9 nm long and 78.9 ± 5.0 nm wide, respectively (Fig. S10C). Notably, partially assembled virus particles in the cytoplasm of the infected cells are also observed. For example, Fig. 7C shows a virus particle in the process of assembly, which has an incomplete coil-stacking structure enveloped by the membrane decorated with spikes and attaches to a piece of coiled nucleoprotein filament to be packaged (indicated by red arrow). These observations suggest that SEV1 completes the process of virion maturation, including envelopment, in the cytoplasm of infected cell.

**Figure 6:**
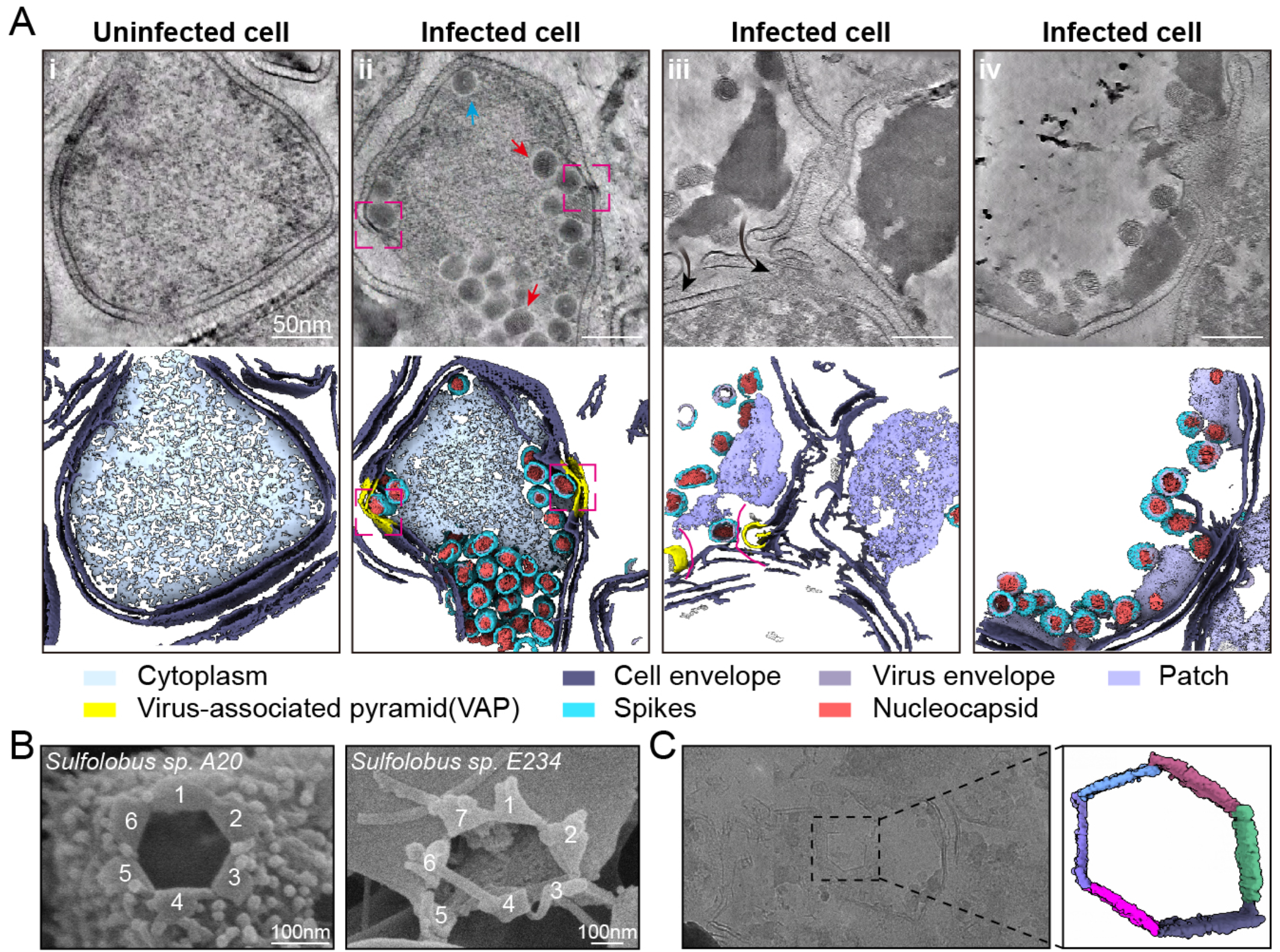
Virion assembly and egress. **(A)** The process of viral assembly and release revealed by cryo-electron tomography (top panels) and segmentations (bottom panels). i. The uninfected cell of *Sulfolobus* sp. A20 with intact cell envelope including the S-layer, cell membrane and ribosomes that are evenly distributed in the cytoplasm. ii. A large number of enveloped viral particles in the cytoplasm, which are arranged in varied orientations (shown by red arrows) with ribosome-like particles surrounding them. VAPs, formed on the cell membrane, are extruding from the S-layer. iii. Patch-like structures in the cytoplasm. Mature viral particles are released from the host cell through the opening VAP. iv. The host cell becomes empty with only a few virus particles and condensed cytoplasmic remnants left. **(B)** The SEM images of opened VAPs in *Sulfolobus* sp. A20 (left panel) and *S. islandicus* E233S overproducing SEV1-ORF84. **(C)** A representative projection at low-magnification and the corresponding segmentation of a VAP in the dashed black box.

**Figure 7:**
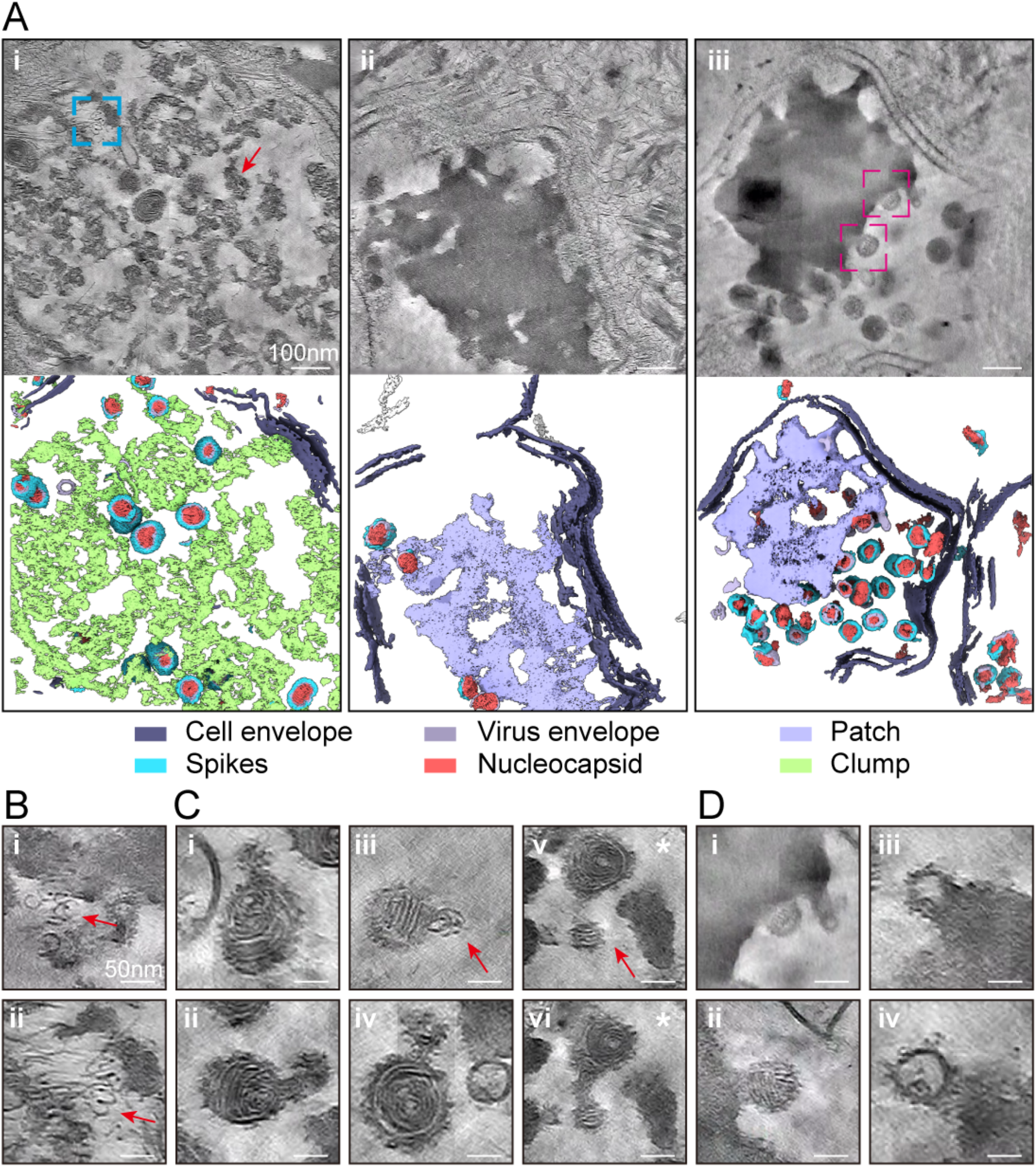
Host cytoplasmic transformation and virus intermediates during SEV1 infection. **(A)** The process of cytoplasm condensation during virion production and assembly, as revealed by tomograms (top panels) and segmentation (bottom panels). **i.** Initial stage of cytoplasmic transformation. The host cytoplasm becomes condensed into small lumps (shown by red arrow). The nucleoprotein filament for virion assembly is highlighted by the blue frame and zoom in B, ii. **ii**. Condensation of the cytoplasm into high density patch-like structures. **iii**. Assembly of virions on the edge of the highly condensed patch-like structures. **(B)** Zoom-in views of the nucleoprotein filaments identified in A. i and other slices. The nucleoprotein filaments undergoing condensation into the lump are indicated by red arrows. **(C)** Virus intermediates identified in the cytoplasm. **(D)** Detachment of nascent virions (complete and incomplete) from the patches (i, ii and iii, iv).

We showed previously that SEV1 exit the host cell through hexagonal virus-associated pyramids (VAPs) on the host cell surface [19]. In this study, we looked closely into this process by using scanning electron microscopy (SEM), cryo-ET and cryo-FIB. In the infected cells, sites on the S-layer where VAPs were to form can be observed (Fig. 6A. ii). The hexagonal aperture of the VAP is clearly shown by SEM (Fig. 6B, left panel). Detailed segmentation reveals that an equilateral hexagon aperture comprises six straight linear segments, with each measuring 100 nm (Fig. 6C). Mature virions and VAPs are observed to co-exist in a host cell, suggesting that the formation of VAPs coincides with the maturation process of the virus (Fig. 6A). A VAP then evolves to form a channel, allowing progeny virions to exit (Fig. 6A. iii, yellow).

Sequence comparison reveals no SEV1 proteins homologous to known VAP proteins [18]. Since the known VAP proteins are all short peptides of ~100 amino acid residues and have a similar fold consisting of an N-terminal transmembrane domain followed by 2-4 α helices [27], we predicted the secondary structure of each protein of SEV1 using the integrative tool PredictProtein [28], and found that ORF84 is a candidate for a VAP protein as it contains an N-terminal transmembrane helix followed by three α helices (Fig. S11). To determine if SEV1-ORF84 is indeed a VAP protein, we cloned its encoding gene under the control of an arabinose promoter in *Sulfolobus islandicus* E233S, which is phylogenetically closely related to *Sulfolobus* sp. A20 and can be readily manipulated genetically [29]. The ORF84 overproducer was grown to the exponential phase under the induction conditions, and the cells were observed under SEM. Opened VAPs with petals are seen on the surface of the cells (Fig. 6B, right panel), indicating that SEV1-ORF84 is a VAP protein in SEV1. Interestingly, VAP appears in a hexagonal form in SEV1-infected *Sulfolobus* sp. A20 but in a seven-folded form in *S. islandicus* E233S producing SEV1-ORF84 (Fig. 6B), suggesting a role for the host cell in VAP assembly.

Our results indicate that the cytoplasm of the host cells undergo considerable changes following the infection of SEV1. Initially, the homogeneous cytoplasm with evenly distributed ribosomes is condensed into small lumps, which are presumably aggregates containing virus components since the density of DNA is clearly visualized around the lumps (Fig. 7A. i). The lumps then continue to condense, forming patch-like structures and creating empty spaces in the cytoplasm (Fig. 7A. ii). Intriguingly, the assembly intermediates of virions appear to emerge from both the patches (Fig. 7A. iii, Fig. 7D) and the surrounding cytoplasmic space (Fig. 7C). Partially assembled virus particles and small-than-normal empty virus envelopes are found on the edge of the condensed cytoplasmic patches, suggesting the active assembly of virions in the patches (Fig. 7D). We speculate that the patch-like structure represents the proposed ‘virus factory’ formed by the host cell in response to virus infection [16, 30, 31], and it continues to function into the late state of infection. Towards the end of the infection cycle, the infected cells become empty with only condensed cytoplasmic remnants and virions left (Fig. 6A. iv). In addition to virus assembly associated with the ‘virus factory’, virus intermediates are also captured in the host cytoplasm (Fig. 7C). Remarkably, these virus intermediates contain the coil-stacking structure and the spike-studded envelope, with a part of the partially assembled nucleoprotein filament hanging outside the particle (indicated by red arrow in Fig. 7C). It is speculated that, as VP4 binds to the viral DNA to form the nucleoprotein filament, the filament would coil into a growing helix until it is constrained by the virus envelope (Fig. 7C). Together, these observations provide insightful details to the intracellular assembly of SEV1 and sub-cellular re-organization in archaeal cell in response to virus infection.

## Discussion

In this study, we show that the SEV1 nucleocapsid consisting of a linear nucleoprotein filament displays a lattice-like appearance, and VP4, the major structural protein of SEV1, is the protein component of the nucleocapsid (Fig. 3A and Table S1). The nucleoprotein filament is organized as helical coils and form a coil-stacking architecture in the SEV1 virion (Fig. 1E and 7B). Based on our data, we propose a coil-stacking model for the nucleocapsid of SEV1 (Fig. 8A). In this model, the viral genomic DNA (Fig. 8A I) is fully covered by VP4 to form nucleoprotein filament with a ‘beads-on-a-string’ appearance, which is subsequently coiled into a spiral structure (Fig. 8A II and S12). VP4 binds the DNA as dimers, which form a channel through which the DNA passes (Fig. 3D). As VP4 binds viral DNA, the resulting nucleoprotein filament coils to form a layer or disc in the spiral structure of the nucleocapsid (Fig. 8A III). Coiling continues for at most five circles (Fig. 8A IV), and the filament will then start coiling in the next layer either from the innermost or outermost coil (Fig. 8A V). A mature virion contains a 16-layer coil-stacked nucleocapsid with a central channel (Fig. 8A VI). The central channel of the nucleocapsid is 11 nm in diameter, suggesting that the innermost coil is 101 bp in length, which is considerably shorter than the persistence length of dsDNA (~150 bp) [32]. Therefore, binding by VP4 strongly bends DNA, permitting the tight packaging of the DNA in a virion.

**Figure 8:**
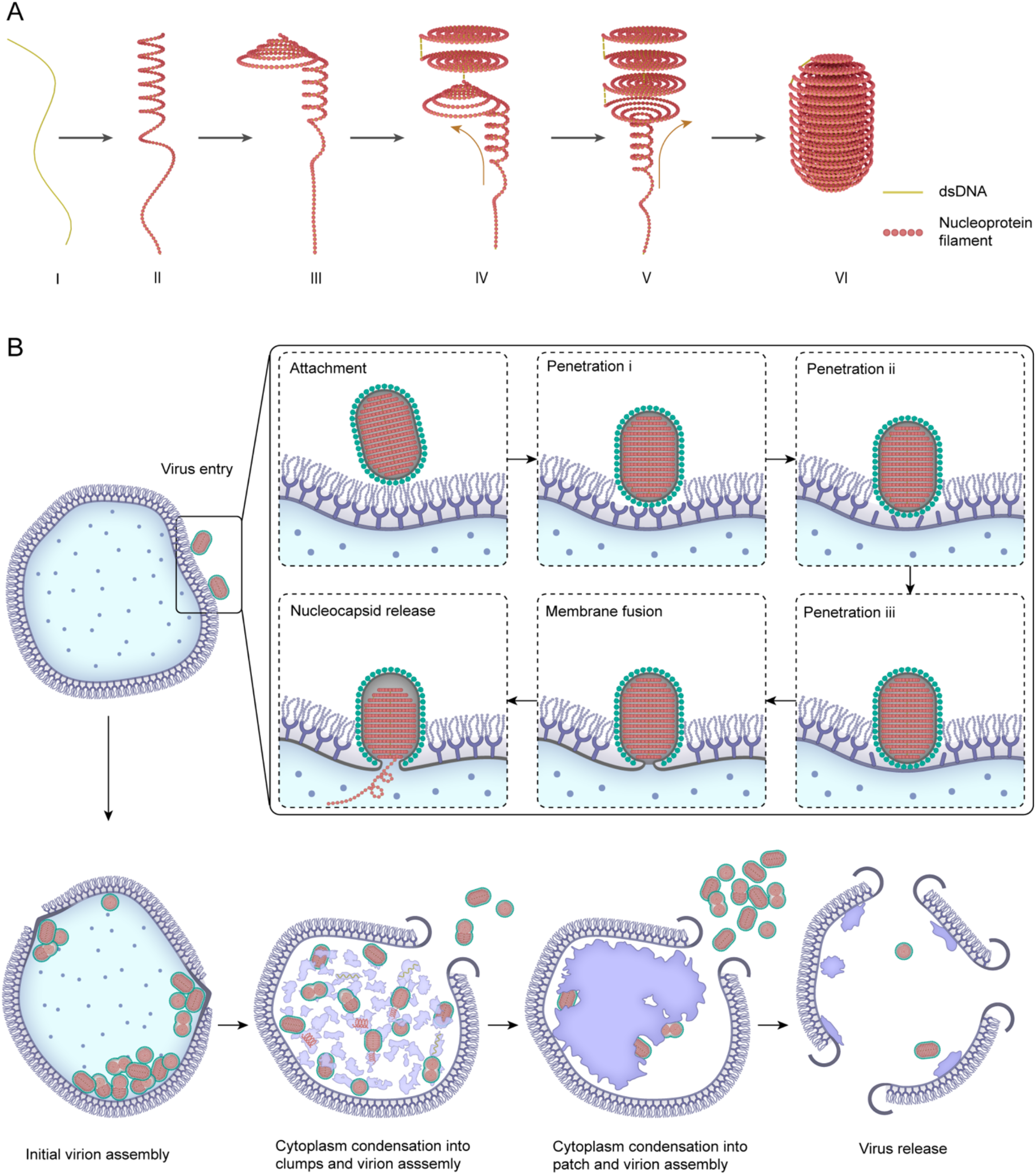
Models of SEV1 assembly and infection. **(A)** A coil-stacking model for SEV1 assembly. **(B)** A diagram showing a cycle of infection by SEV1.

The SEV1 nucleocapsid is organized like a spool, somewhat analogous to the SPV1 nucleocapsid in the general appearance, although the nucleocapsids of the two viruses are assembled in distinctly different manners [3, 18]. In SEV1, the nucleoprotein filament first coils into a disc-like structure, and the disc-like structures stack into the nucleocapsid. By comparison, in SPV1, the nucleoprotein filament forms rafts-like structures and the “rafts” spool to form the nucleocapsid [3]. Despite the differences, however, the nucleoprotein filaments of the two viruses are probably organized by their respective structural protein in a similar manner [3]. Notably, VP4 from SEV1 displays no significant sequence to VP1, the MCP of SPV1, indicating that the similar genome packaging strategies employed by the two viruses were evolved through distinct pathways.

In addition to SEV and SPV1, several other archaeal viruses from extreme habitats, such as rudivirus SIRV2, lipothrixvirus SIFV and tristromavirus PFV2, also display the ‘beads-on-a-string’ structure [6]. SEV1-VP4 has no sequence similarity to MCPs from these viruses. It is known that SEV1-VP4 is enriched with the basic amino acid residues arginine (R), lysine (K) and asparagine (N), which form ionic contacts with DNA. RKN account for 20% of the total amino acid residues in SEV1-VP4. Similar RKN enrichment is also found in the MCPs from SIRV2 (19%), AFV1 (18 and 19%), SFV1 (17 and 21%), and SPV1 (22%) [3].

MCPs from these viruses have the following common features. (1) They are all small proteins ranging in size from 78 to 204 amino acid residues. (2) They have similar overall folds consisting of five helices for DNA binding. (3) They are enriched in RKN amino acids for interaction with DNA. (4) They bind DNA in the form of a dimer, creating a channel that allows the DNA to pass through. These shared features suggest that the genomes of these archaeal viruses, which are not closely related, may be packaged in a similar fashion with the ‘bead-on-a-string’ characteristics, and this DNA packaging strategy may represent an adaptation of archaeal viruses to living in extreme habitats.

Phylogenetic analysis reveals that the archaeal viruses with SEV1-VP4 homologues are widely distributed in acidic and thermal habitats across the Pacific Rim. The distribution of SEV1-VP4 homologues may be even more extensive as it might be underestimated due to the limitation of sampling bias and sequencing depth. However, the geographic distribution of these homologues does not appear to correlate with their phylogenetic distribution. While the SEV1-VP4 homologues share low amino acid sequence similarity (34-47%) (Table S2), the predicted 3-D structures of these proteins exhibit high structural similarity (Fig. S5C). These results point to the possibility that the high variation of SEV1-VP4 homologues at protein sequence level and conservation at 3-D level might be a result of adaptive evolution in response to the acidic and thermal environment challenges, rather than merely a consequence of spatial distribution and gene flow.

Archaeal viruses infecting extremophiles exhibit extraordinary structural diversity. To date, available structures indicate that most of these viruses, including rudivirus SIRV2 [7], lipothrixivirus SIFV and AFV1, and tristromavirus PFV2 [6], exist as rigid and compact particles, whose robustness is suggested to be a critical factor in their adaptability to extreme environments [4]. However, some viruses with flexible and fragile structures, such as SSV19, STSV2 and His1 [5, 33, 34] as well as HRPVs [12], also thrive in extreme environments. The flexibility of SEV1 virion appears even more pronounced as irregular virions with one half of the nucleocapsid flipping 90° or 180° exist (Fig. 2). This degree of flexibility in nucleocapsid is rarely seen in other viruses. Comparing to the rigid viruses, the significantly high flexibility and diversely different morphs of SEV1 might help to keep its stability, possibly contributed by the local stable coil-stacking pattern formed by VP4 under the extreme environments.

The morphological changes of intact and naked SEV1 virions under different conditions provide further insights into SEV1 stability in extreme environment. It was reported that dsDNA can remain soluble in pH 5-8 but precipitate in acidic solutions and degrade in alkaline solutions [35]. Our results indicate that the SEV1 virion and its nucleocapsid remain stable under the harsh conditions of pH 2-5. This finding implies that VP4 substantially stabilizes the nucleocapsid. The increased pH stability of the nucleocapsid is consistent with its life cycle during which it undergoes sharp pH changes as the virion is assembled in the cytoplasm (pH 6.5) [36] and released into the hot spring water (pH 2-3). The virus envelope is damaged above pH7 (Fig. 4A) and the nucleocapsid is completely disrupted above pH11 (Fig. 4A). On the other hand, the intact SEV1 virion is morphologically stable at high temperature (100°C) (Fig. 4A) but the unenveloped nucleocapsid is not (Fig. 4B). We conclude from these results that the lipid envelope is crucial for the thermostability of the SEV1 virion but not critical for the pH stability of the nucleocapsid.

The lipid envelope of SEV1 is composed of GDGT-0, 1, 2, 3, 4, with GDGT-4 being the most abundant composition [18]. The inclusion of an increasing number of cyclopentane rings leads to tighter packing of the molecules, thus increasing the thermostability of the lipid membrane and reducing permeability [37], especially to small ions [38]. SEV1 is not protected by a capsid shell and, instead, maintains the structural integrity through the increased rigidity of the lipid envelope. Hence, the enrichment of GDGT-4 in the lipid envelope may suggest an adaption to the highly flexible architecture of the virus at elevated temperatures. Moreover, it reduces permeability to protons, contributing to the stability of SEV1 in extremely acidic environments [39].

We show that SEV1 initiates entry by attaching first to the densely present glycan chains on the S-layer of the host cell (Fig. 5A). VP1, VP2 and VP3 from SEV1 are predicted to function as hydrolase based on a protein language model [25], and VP1 and VP2 are presumably attached to the virus envelope via their transmembrane domains [18]. Therefore, we speculate that these viral proteins contribute to host recognition and penetration by the virion (Table S2). The disruption of the S-layer of the host cell following its interaction with SEV1 is consistent with the speculation (Fig. 5B). S-layer disruption presumably allows the virus to interact with the host membrane, initiating subsequent membrane fusion (Fig. 5C). This is distinct from the entry model of HRPV-6, where membrane fusion mediated by the extended conformation of virus spikes triggered by heat treatment [12] [40]. In contrast, SEV1 spike shows no changes under the applied infection conditions (pH3, 75°C), even the increasing temperatures up to 100°C (Fig. 4A, last panel). Instead, the conformational changes in the SEV1 spikes are triggered by a decrease in pH from 2 to 0.5 (Fig. 4C), suggesting that extended conformation of the spike may not be the key factor for the SEV1 entry. This implies that the mechanisms mediating membrane fusion may differ among archaeal viruses which remain to be explored.

The infection cycle of archaeal viruses has been investigated [17] [30] [31]. However, the knowledge of steps during virion assembly in a host cell is rudimentary. In this study, we observed a series of viral assembly intermediates, including unpackaged nucleoprotein filaments, partially assembled nucleocapsid with short filaments hanging outside, small membrane vesicles without genomic contents, etc., in the host cells infected with SEV1. We also found that the host cytoplasm underwent visible condensation to form high density patches with the SEV1 virion particles and assembly intermediates at the edge (Fig. 7A iii). Although the exact nature of the patches is unclear, the possibility exists that they result from the coupling between viral DNA replication and viral protein synthesis as well as the coupling between the synthesis and assembly of virions, thus serving as virus factories. Moreover, these virus factories produced nascent virions successively till the late stage of infection, during which the cell became empty. This may represent a strategy for SEV1 to replicate the virions efficiently under the harsh environmental conditions.

Virus egress by VAPs is unique in archaea and adopted by archaeal viruses with different morphologies, particularly the unenveloped archaeal viruses, i.e. icosahedral virus [17], filamentous viruses along with their relatives [16, 30]. Conventionally, enveloped archaeal viruses, e.g., SSV1 and STSV2, are known to acquire the lipid membrane during the process of budding [14]. The endosomal sorting complexes required for transport (ESCRT) machinery used by eukaryotic virus egress [41] play roles in the egress of STSV2 [42]. We showed previously that intact SEV1 virions with the envelope are formed inside the infected cell and exited the host cell through VAPs [18]. We have now identified the VAP protein of SEV1. The SEV1-VAP appears to share similar folds with those from other archaeal viruses, despite their dissimilarity in amino acid sequence, supporting the notion that the VAP-based egress pathway is an ancient strategy of virus release.

Based on the results of this and previous studies, we propose a model for the infection cycle of SEV1 (Fig. 8B). An SEV1 virion initially recognizes and attaches to glycan chains on the S-layer of the host cell, and then disrupts the S-layer to get in touch with the cell membrane. Membrane fusion occurs to allow the release of SEV1 nucleocapsid into cytoplasm of the host cell. Viral DNA replication and protein synthesis then take place. The host cytoplasm undergoes condensation into lumps and further into patch-like virus factories. The progeny virions are assembled in the cytoplasm and, simultaneously, VAPs formed on the cell surface. Finally, VAPs open to allow the release of the progeny virions, leaving behind an empty host cell.

## Methods

### Growth conditions of the host strain and the virus

The virus *Sulfolobus* ellipsoid virus 1 (SEV1) was isolated from the acidic hot spring of Costa Rica, together with its host strain *Sulfolobus sp*. A20 [18]. *Sulfolobus* sp. A20 was grown in the Zillig medium at 75 °C with shaking at 150 rpm [43]. For virus production, a sample (1 ml) *Sulfolobus* sp. A20 grown to OD_600_ of 0.2-0.4 was mixed with an SEV1 virus lysate (200 µl) at 75°C. The mixture was inoculated in the 250-ml fresh medium and incubated for 7 days as described above. Virus particles were harvested by centrifugation at 15,428 x g for 20 min at 4 °C (F14-6×250y, Thermo). The virus-infected culture was cultivated to an OD6_00_ of 1.0 for sample preparation for cryoFIB and SEM described below.

### Virus-cell fusion assay

SEV1 virions (4 mg/ml) were mixed with R18 dye in the DMSO to final concentrations of 1 µM dye and 0.1% DMSO. After incubation at 37°C for 1 h, the free dye was removed by G25 desalting column (GE healthcare). The host cells were grown to an OD_600_ of 0.2 and collected by centrifugation at 8000 x g for 15 min at 25°C. The cells were resuspended in Zillig’s basal salt solution. The host cell suspension was mixed with R18-labelled SEV1 virions for 1.5 h at 75°C or 23°C. Samples were observed under the Leica TCS SP8 STED microscope fitted with a 100 X oil objective lens, and images were processed by Leica Application Suite X.

### Purification of virus particles and sample preparation for electron microscopy

The SEV1 virions were collected by Amicon stirred cell (400 mL, Merck) and purified by CsCl density gradient ultracentrifugation (214,200 x g for 8 hr at 4°C). Discontinuous gradients of CsCl were set up as 0.3, 0.4, 0.5 and 0.6 g/ml. The fraction containing virus particles, located at ~0.4 mg/ml, was collected by using a syringe, and subsequently dialyzed against Zillig’s basal salt. For cryo-EM analysis, 3 µl of the sample was applied to NiTi grids, which were glow-discharged for 120 sec, then the samples were blotted and vitrificated by plunging into liquid ethane with a Vitrobot (FEI) operated at 4°C and 100% humidity.

### Determination of the nucleoprotein of SEV1

To determine the nucleoprotein of SEV1, we extracted its nucleoprotein filament as follows. The purified SEV1 virions were treated with 1% NP-40 for 30 min, and the treated sample was centrifuged at 3,000 x g for 10 min at 4°C. The pellet and supernatant were subjected to 15% SDS-PAGE together with the purified SEV1 virions. The gel was stained with Coomassie brilliant blue G250. Only one band was visible from the pellet and the band was subjected to liquid chromatography-tandem mass spectrometry (LC-MS/MS). For MNase digestion, the purified SEV1 virions were dialyzed against ddH_2_O, and treated with 18 cycles of freeze (−80°C) and thawing (25°C) to release the nucleoprotein filaments [3]. The resulting sample was digested overnight at 37°C with MNase (2000 U, NEB), and subjected to 1% agarose gel electrophoresis.

### Heterologous expression and SEM analysis of SEV1-ORF84

SEV1-ORF84 was amplified by PCR from the SEV1 DNA [18]. The gene sequence was inserted into pSeSD plasmid under the control of an arabinose-inducible promoter [44]. The resulting construct was introduced into *Sulfolobus islandicus* E233S by electroporation [29]. The transformed cells were cultured in modified Zillig’s medium supplemented with 0.1% tryptone [43]. The single colony of the transformed cells was selected. After two passages, the culture was grown to an OD_600_ of 0.6, and arabinose was added. Following the growth of the culture to an OD_600_ of 1.0, cells were collected for scanning electron microscopy (SEM) [18].

### Stability of SEV1 under pH and high temperature

To investigate the stability of SEV1 under extensive pH fluctuation, aliquots of 10 µL virus preparation were mixed with equal volume of HCl solution at pH0.5, 1, 2, 3, 5, 7, 9, 11, 13. The mixtures were kept at room temperature for 30 min. The thermostability of SEV1 was explored at 45 °C, 55 °C, 65°C, 75 °C, 85 °C and 100 °C (boiling water bath). Each aliquot of 10 µL virus preparation was treated at the above temperatures for 30 min. To obtain naked SEV1 nucleocapsid, the purified SEV1 virions were treated with 0.2% Triton X-100 for 30 min at 25°C. The detergent was removed by membrane dialysis (0.025µm MCE membrane, MF-Millipore) against Zillig’s basal salt for 30 min.

After the treatments, for the virions treated at temperature of 100°C, a modified vitrification procedure and apparatus with an additional centrifuge tube adapted from reference [22] was applied to ensure that viruses are maintained at high temperatures at the moment of freezing. In addition to the vitrification apparatus, filter paper grids and pipette tips were kept at a temperature of 60°C as much as possible. The 7 µl suspension was applied to NiTi grids which were glow-discharged for 120 sec, then the samples were blotted and vitrificated by plunging into liquid ethane with a Vitrobot (FEI) operated at 60 °C and 100% humidity. The parameters of wait time, blot force and blot time is 1s, 0N and 1.5s. For the virions treated at other conditions, the 3 µl high-temperature suspension was applied to NiTi grids which were glow-discharged for 120 sec, then the samples were blotted and vitrificated by plunging into liquid ethane with a Vitrobot (FEI) operated at 4 °C and 100% humidity.

### Cell infection and vitrification

To explore the virus entry penetrating the S-layer, SEV1 virions (2mg/mL) were added to the logarithmic phase culture of (OD_600_~0.2) of *Sulfolobus sp*. A20 to reach the superinfection to observe the process of the virus entry by CryoET. The samples were incubated at 75°C for 5min, 10min and 30min and prepared to vitrify. To prevent the two-side filter paper blotting and the low temperature which could result in the failure to capture the entry, one-side blotting method was used for sample freezing. The high-temperature mixture of cell and SEV1 was immediately applied to NiTi grids (2/1) which were glow-discharged for 120 sec. And then blotted for 1.5 seconds by Leica GP1 (Leica Microsystems), followed by plunge freezing in liquid ethane.

The virus-infected culture was set up as described in the section of “Growth conditions of the host strain and the virus”. 30mL of the infected culture was cultivated to OD_600_∼1 and harvested by centrifugation at 8000 x g for 15 min at 25 °C. The cells were resuspended by 1mL Zillig’s basal salt and an aliquot of 3 µl cell suspension with different MOI was applied to the glow-discharged holey carbon-coated copper (R 2/1, 200 mesh) (Quantifoil) and blotted for 9 seconds by Leica GP1 (Leica Microsystems), followed by plunge freezing in liquid ethane.

### Cryo-lamella preparation using cryo-FIB

Vitrified cells were further processed by cryo-FIB milling for the preparation of lamellae. A dual-beam microscope FIB/SEM Aquilos 2 (Thermo Fisher Scientific) equipped with a cryo-transfer system (Thermo Fisher Scientific) and rotatable cryo-stage cooled at −191°C by an open nitrogen circuit was used to carry out the thinning. During the cryo-FIB milling process, the milling angle between the FIB and the specimen surface was set to 5-10°. The milling was performed parallel from two sides to produce vitrified cell lamella [45]. The accelerating voltage of the ion beam was kept at 30 kV, and the ion currents were in the range from 0.43 nA to 40 pA. The rough milling utilized a strong ion beam current of 0.43 nA and the final fine milling was operated with a small ion beam current of 40 pA. The thickness of the residual thin lamella with a good quality was < 150 nm.

### Acquisition and processing of cryo-ET tilt series

The tilt series of purified SEV1 virions were collected with Titan Krios G3 (Thermofisher Scientific) 300 KV TEM, equipped with a Gatan K2 direct electron detector (DED) and a BioQuantum energy filter. Tilt series were collected under a magnification of ×105,000, resulted in a physical pixel size of 1.36 Å, under counting mode, in K2 DED. The total dose was set to 3 electrons per square angstrom per tilt, fractioned to 10 frames in a 1.2 s exposure, and the tilt range were set to be between –55° to +55° with a 3° step, resulting in 35 tilts and 115 electrons per tilt series. The slit width was set to be 20 eV, with the refinement of zero-loss peak after collection of each tilt series, and nominal defocus was set to be –3.5 to –4.5 µm. Tilt series of SEV1 virions treated with different temperature and pH conditions were collected with the same parameters as above except using a physical pixel size of 1.76 Å.

For the cryo-FIB milling sample, before data collection, the pre-tilt of the sample was determined visually, and the pre-tilt was set to be 10° or –9° to match the pre-determined geometry caused by loading grids. And the tilt range were set to be between –60° to +40° for –10° pre-tilt or –40 to +60° for +10° pre-tilt, with a 3° step, resulting in 33 tilts and 99 electrons per tilt series. The rest of the collection details are consistent with those mentioned above. All tilt series used in this study were collected using a dose-symmetry strategy-based beam-image-shift facilitated acquisition scheme, by in-house developed scripts within SerialEM software [46-48].

All the tilt movies were processed in Warp [49] to generate the stacks and then aligned using IMOD [50]. The reconstructions were performed using SIRT-like filter and binned 4 times implemented in IMOD [50]. The missing wedge of all reconstructions were recovered in IsoNet [51].

### Tomogram Visualization and Segmentation

Segmentations shown were generated using Dragonfly (v.2022.2) [26]. For the purified SEV1 tomograms, a 5-class U-Net with 2D input was trained on 5 tomogram slices to detect background voxels, spike, envelope, bacterial flagella and genomic contents. And for the intracellular tomograms, the multi-ROI U-Net was trained to detected cell membrane, virions (spike, envelope, and genomic contents) and cytoplasmic content. All segmentations produced by the neural network were cleaned up in Dragonfly, subsequently exported as binary tiff and converted to mrc files using ‘tif2mrc’ in IMOD. All segmentations were visualized using ChimeraX 1.3 [52].

### Subtomogram Averaging and Classification

The tomograms were binned 8 and subjected to the N2N [53] method to denoise in order to facilitate the particle picking. For the NPs structure which was shown in Fig. 2C green box, particle selection was conducted by pairing two NPs within the same disc layer, and the center positioned middle between the two NPs. The 500 coordinates were manually picked in Dynamo [54], then transferred back to Warp for exporting of sub-tomograms in box size of 60^3^ voxels with pixel size of 2.72 Å. One random subtomogram was used as the initial model. 3D classification was then performed in Relion (T=1) to obtain a smooth averaged density [55, 56]. Finally, six NPs were identified in the averaged density map.

For the NPs structure which was shown in Fig. 2C orange box, particle selection was ocating the box center between two regular discs. The 246 coordinates were manually picked and then transferred back to Warp for exporting of sub-tomograms in box size of 70^3^ voxels with pixel size of 2.72 Å. One random subtomogram was used as the initial model. 3D classification was then performed in Relion (T=1) to obtain a smooth density. Finally 18 NPs were identified in the density map.

For the spike structure which was shown in Fig. 2E, we manually picked the enveloped point and the middle of spike points to give two of three Euler angles in Dynamo. And then we restricted two Euler angles (--sigma_tilt and --sigma_psi in RELION) to 3D classification.

For the structure of genomic content in Fig. 2C purple box, we created a dipoleSet model in Dynamo to annotate the centres and edge points of SEV1 virions, and then converted these dipoleSet models into oversampled vesicles with a function which used in-house script. The spacing we sampled was 10nm and each particle containing spike, envelope and genomic content was obtained an initial orientation in order to facilitate subsequent alignment. The orientation tbl file containing 7000 particles was then transferred back to Warp for exporting of sub-tomograms in box size of 100^3^ voxels with pixel size of 5.44 Å. Afterwards, we employed masks of varying sizes (35nm, 40nm, 45nm, 50nm) for local alignment and classification, observing discrepancies in the classification outcomes. All averaged results were then consolidated for subsequent genomic content analysis.

Sub-tomogram refinement was done in RELION 3.0 or 3.1 [55, 56], transform of RELION’s star file and Dynamo’s table file was done by ABTT package [57]. Mask generation was done by combination of Dynamo and/or RELION.

### Phylogenetic analysis of VP4

The protein sequence of SEV1-VP4 (YP_009639281.1) was submitted to Genbank to search for the similar sequences by psi-BLAST. However, only two sequences were identified including SEV1-VP4 itself. The further attempt was executed by psi-BLAST [58] hitting the database of IMG/VR V4.1 database and 50 sequences were obtained with the parameters ‘-num_iterations 3’ and E-value <10^−5^. The sequences were aligned using MAFFT [60] with the ‘--auto’ parameter. The ModelFinder [61] was used for searching the maximum likelihood phylogenetic models with the following parameters: ‘-madd LG+C20~C60, WAG+C20~C60, Q.pfam+C20~C60 -mrate I,G,R,I+G -mfreq F,FO’. The BIC and AICc criteria were used to select the most fitting model which was ‘LG+FO+G4’. The maximum likelihood phylogenetic analysis was inferred using IQ-TREE2 [62, 63] with parameters ‘-wbtl -bb 1000’.

Sequence alignment between SEV1-VP4 and SBV-GP06 was performed by ClustalW [64] and the results was displayed by ESPript 3.0 [65]. The alpha-helices of the two proteins predicted by AlphaFold described below were also showed by ESPript.

### AlphaFold Structure Prediction

Structural prediction of proteins and protein-DNA complexes were carried out with AlphaFold v2 [66] and AlphaFold v3 [67], respectively. Visualization, figure generation and model docking were performed in UCSF ChimeraX v1.3 [52]. Measurement of the long and short axes of the SEV1 virion was conducted manually using the measurement tool in IMOD.

## Supporting information

Supplemental Table 2. Sequence information of SEV1-VP4 homologues

## Data availability

The sub-tomogram averaged cryo-EM maps, including the average map of six beads composed of the nucleocapsid protein from SEV1(EMD-62178), the average map of eighteen beads composed of the NP (EMD-63241), the lattice-like arrangement of nucleocapsid beads (EMD-62348), the DNA coil forming by the nucleoprotein filaments in (EMD-62342), and the averaged map of pronounced stacking discs with a conspicuous channel (EMD-62349), have been deposited in the Electron Microscopy DataBank (EMDB), and will be available upon publication or by reviewer request. Any additional information required to reanalyze the data reported in this paper is available from the lead contact upon request.

## Acknowledgements

This work was supported by grants from the National Natural Science Foundation of China (grant no. 41902316, 92351001, 32241029, 92051109), the Chinese Ministry of Science and Technology (2023YFA0913400, 2021YFA1300100), the PI Project of Southern Marine Science and Engineering Guangdong Laboratory (Guangzhou) (GML20240002), the Chinese Academy of Sciences (CAS) (XDB3700000, JZHKYPT-2021-05) and the China Postdoctoral Science Foundation (grant no. 2018M640159). All EM data were collected and processed at the Center for Bio-imaging (CBI), Institute of Biophysics (IBP), CAS. We would like to thank Xujing Li, Lulu Qin and Xiaojun Huang for their technical help and support with electron microscopy and cryo-FIB. We thank Weiwei Niu, Chengcai Pan and Niannian Ding for their assistant in experiment. We thank Dr. Jingfang Liu (Institutional Center for Shared Technologies and Facilities of Institute of Microbiology, CAS), for their work of identification of protein with Mass Spectrometry in this manuscript. We also would like to thank Prof. Meng Li of Shenzhen University and Geng Wu for their valuable discussion.

## Author contributions

P.Z., L.H., H.N.W. and H.N.Z. initiated the project. H.N.Z. and H.N.W. designed the research. H.N.W., H.N.Z., Y.X.F., and Z.F.Z. supplied the experimental materials. H.N.Z., H.N.W., Y.L. and K.S. performed the cryo-EM and cryo-FIB sample preparation. H.N.Z., Y.L. and H.N.W. perform data collection. H.N.Z., H.N.W. analyzed the data. H.N.Z., H.N.W and H.Y.C. contributed to the phylogenetic analysis. H.N.Z., H.N.W., L.H. and P.Z. wrote the paper.

