## Supplemental figures and tables for "Insights into the spool-like architecture and infection strategy of the archaeal virus SEV1"

**Figure S1.** Segmentation of SEV1 virions
**Figure S2.** The slices of SEV1 cryo-ET showing irregular SEV1 virions with partially rotated nucleocapsid in varying orientations
**Figure S3.** Connections between adjacent coiled discs.
**Figure S4.** dsDNA fully covered by VP4.
**Figure S5.** Phylogeny and distribution of VP4
**Figure S6.** Sequence alignment of SEV1-VP4 and SBV-GP06
**Figure S7.** Slices of SEV1 virions treated and prepared under the extreme conditions
**Figure S8.** The structural changes of the naked SEV1 virions with fluctuated temperatures and pH
**Figure S9.** Glycan chains of the S-layer
**Figure S10.** The pipeline of cryo-FIB and the statistic of diameters of SEV1
**Figure S11.** Comparison of secondary structures of PVAPs.
**Figure S12.** Structural elements of the SEV1 nucleocapsid

**Table S1.** Mass spectrometry and MaxQuant generated iBAQ value
**Table S2.** Sequence information of SEV1-VP4 homologues
**Table S3.** Functional annotation of viral proteins by NetGo

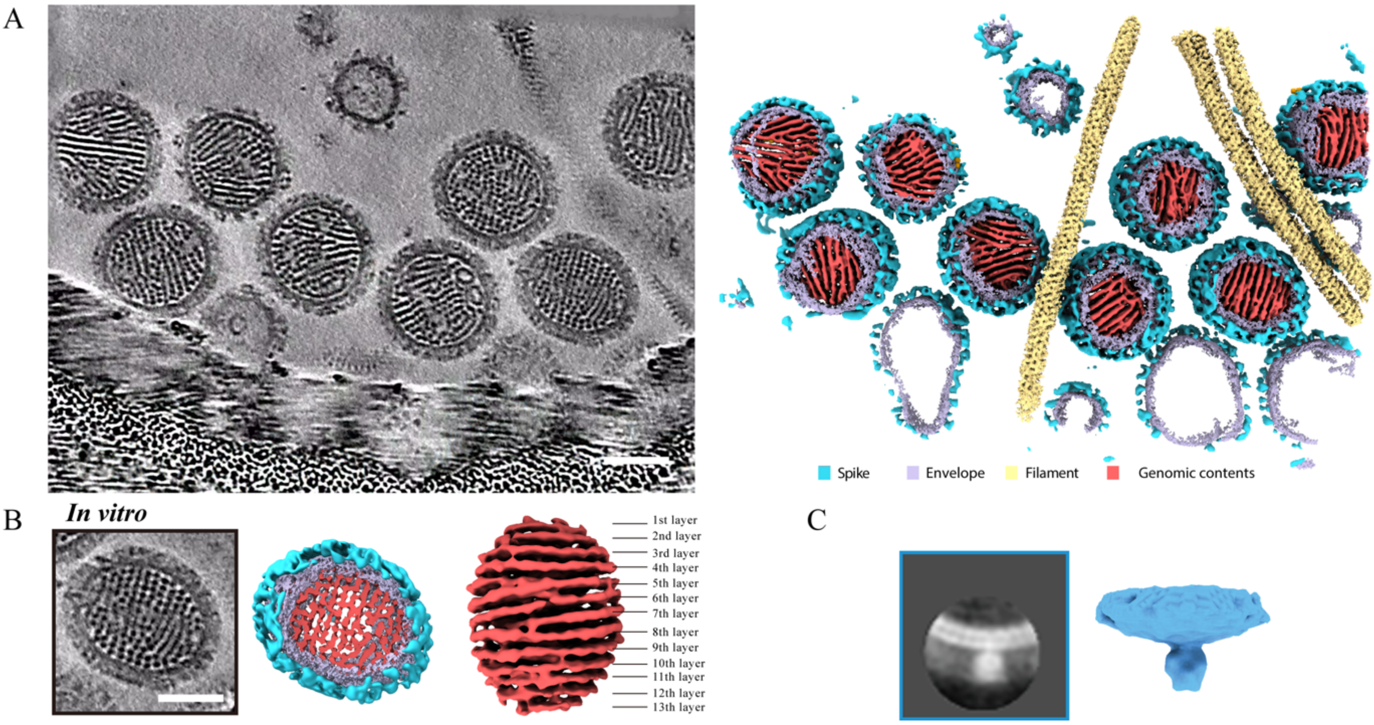


**Figure S1: Segmentation of SEV1 virions.** The colour scheme is showed in the figure (A). **(A)** A slice of cryo-ET of purified virions (left panel) and its corresponding segmentation map (right panel). **(B)** The slice (left panel) and segmentation (middle panel) of a representative SEV1 virion. The right panel displays the multilayered genome content of the SEV1 virion. The scale bar is 50 nm. **(C)** The averaged map of a single spike. The left panel is a slice of the averaged result.

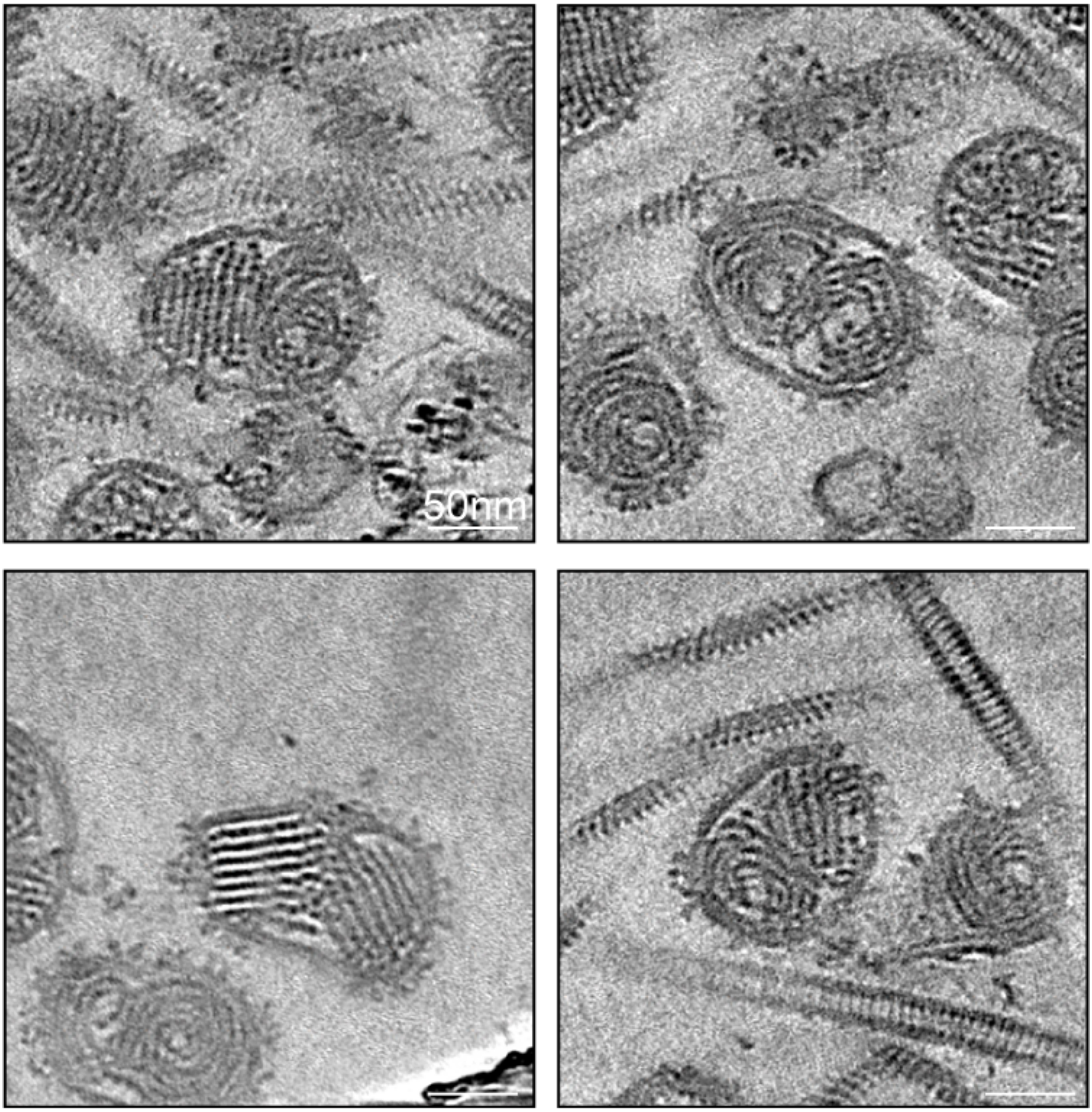


Figure. S2: The slices of SEV1 cryo-ET showing irregular SEV1 virions with partially rotated nucleocapsid in varying orientations.

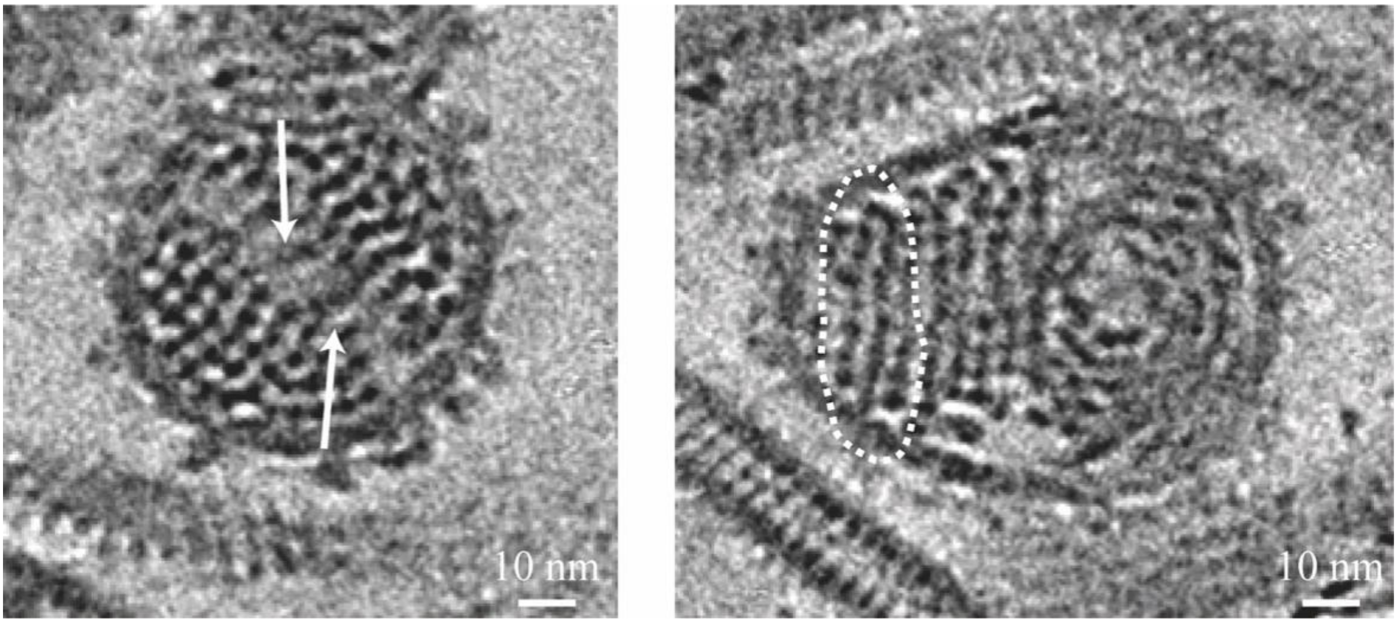


**Figure S3: Connections between adjacent coiled discs.** In the left panel, the white arrows point the linear connection linking the adjacent coiled discs in the interior of the nucleocapsid. In the right panel, the white dashed circle highlights the connections between adjacent discs along the edge of the nucleocapsid.

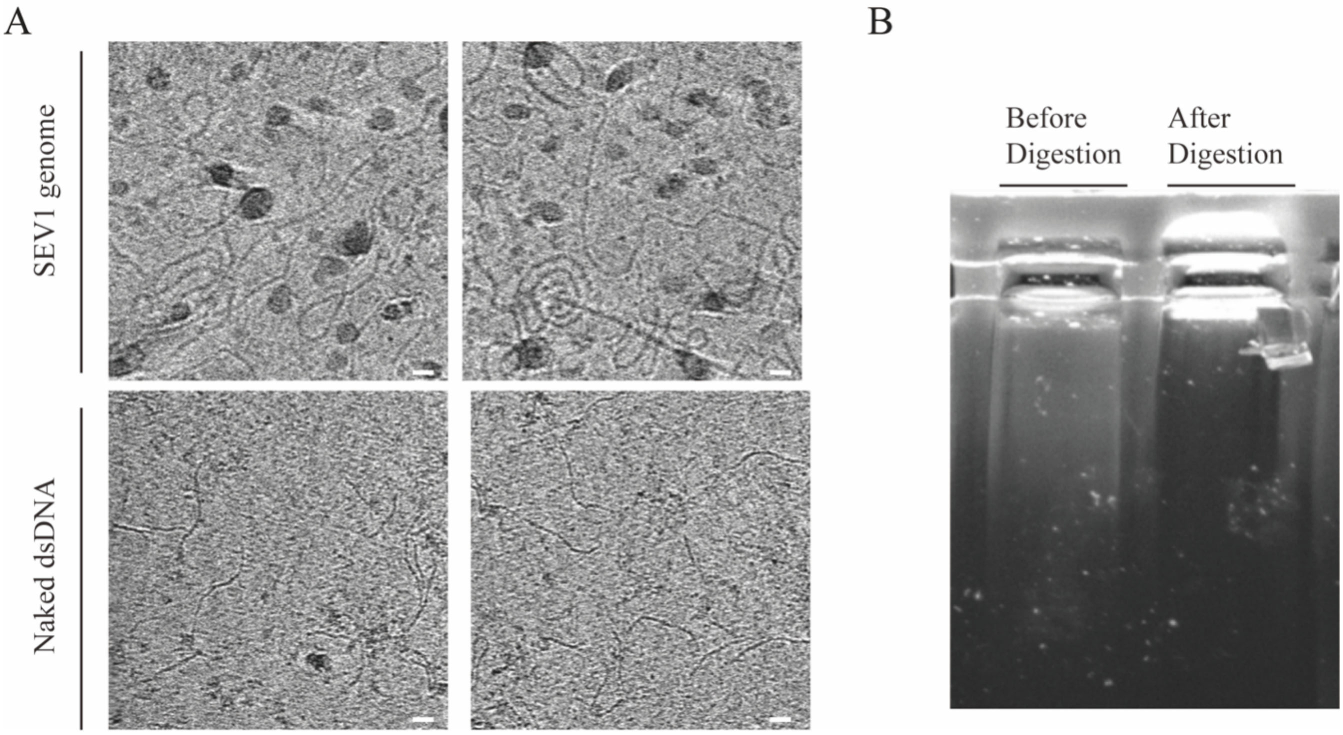


**Figure S4: dsDNA fully covered by VP4. (A)** The slices of cryo-ET of SEV1 genome and naked dsDNA (147bp). The scale bar is 50 nm. **(B)** 1% agarose gel electrophoresis image of nucleoprotein filaments of SEV1 before and after MNase digestion. A large proportion of DNA did not enter the gel.

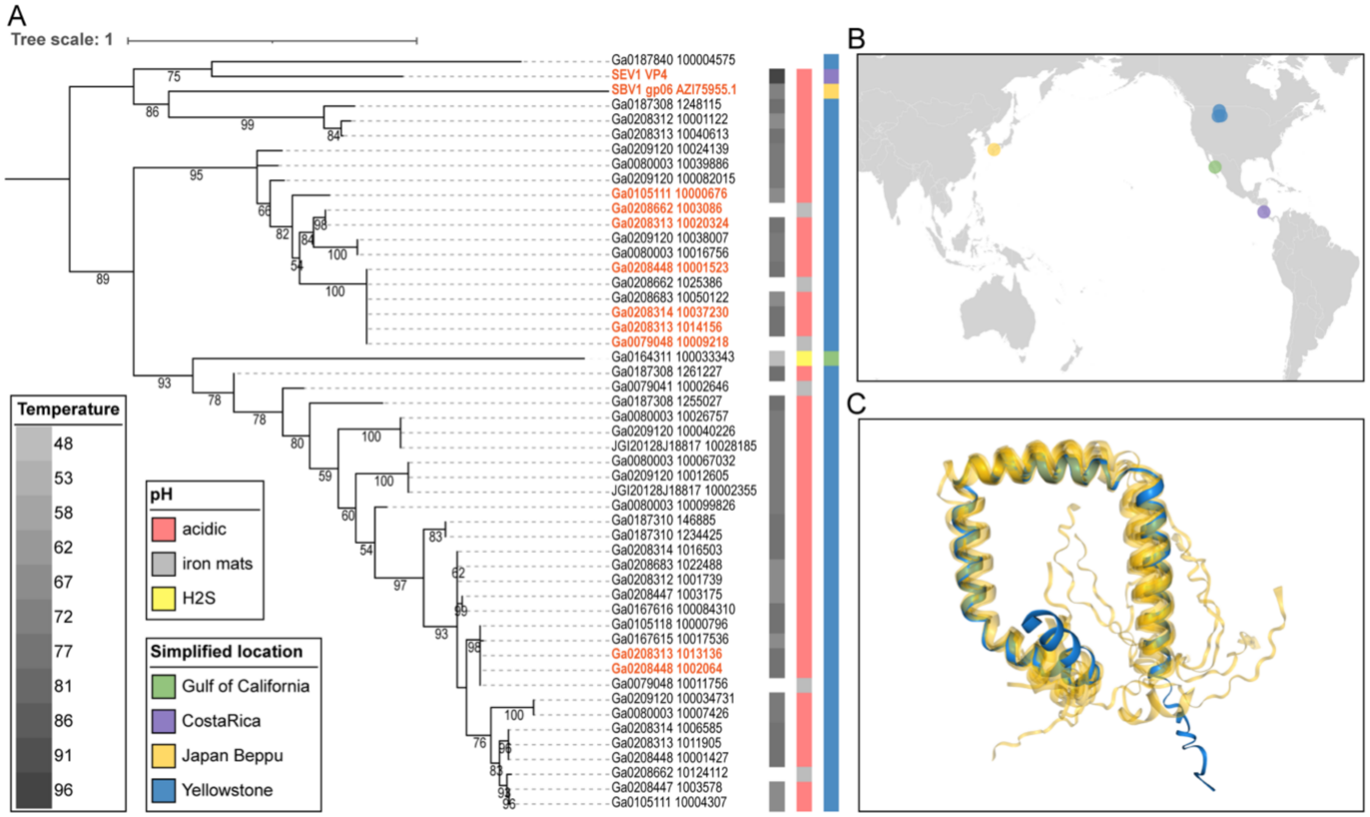


**Figure S5: Phylogeny and distribution of VP4. (A)** The phylogenetic tree of VP4 homologues constructed by maximum likelihood. The available parameters of habitats including sample sites, pH and temperature are listed in the figure. **(B)** The map of the VP4 distribution across the Pacific Rim. The color scheme corresponds to that in (A). **(C)** Alignment of predicted 3-D structures of VP4 homologues. The predicted structure of SEV1-VP4 is highlighted in blue while others are displayed in yellow. The aligned structures are predicted based on the sequences highlight in red in (A)

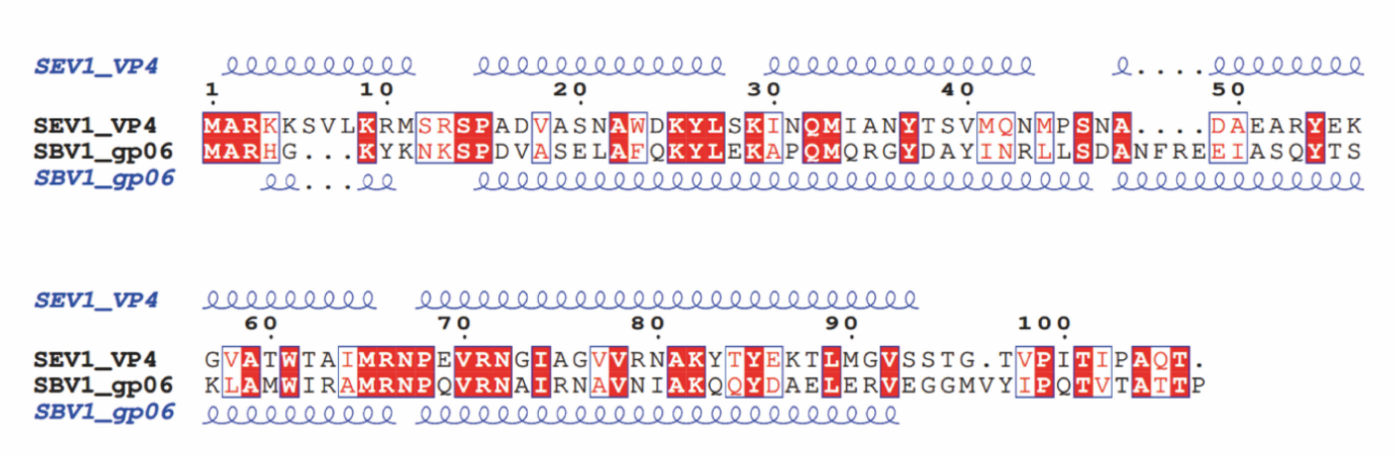


**Figure S6: Sequence alignment of SEV1-VP4 and SBV-GP06.** The predicted alpha-helices are displayed on top (SEV1-VP4) and bottom (SBV-GP06) of the alignment respectively. The identical amino acids between the two aligned sequences are shaded in red backgrounds. Amino acids showed by red letters and enclosed in blue boxes are of similar biochemical characteristics.

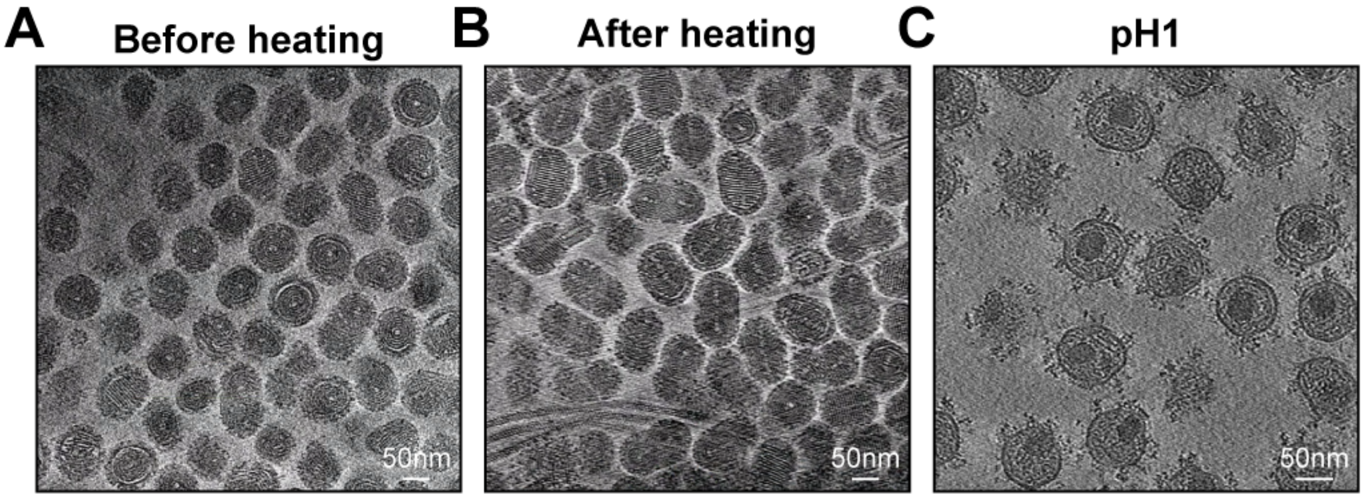


**Figure. S7: Slices of SEV1 virions treated and prepared under the extreme conditions. (A)** SEV1 specimen prepared by regular method. **(B)** The SEV1 specimen treated and prepared at 85°C. **(C)** The SEV1 specimen treated and prepared at pH1.

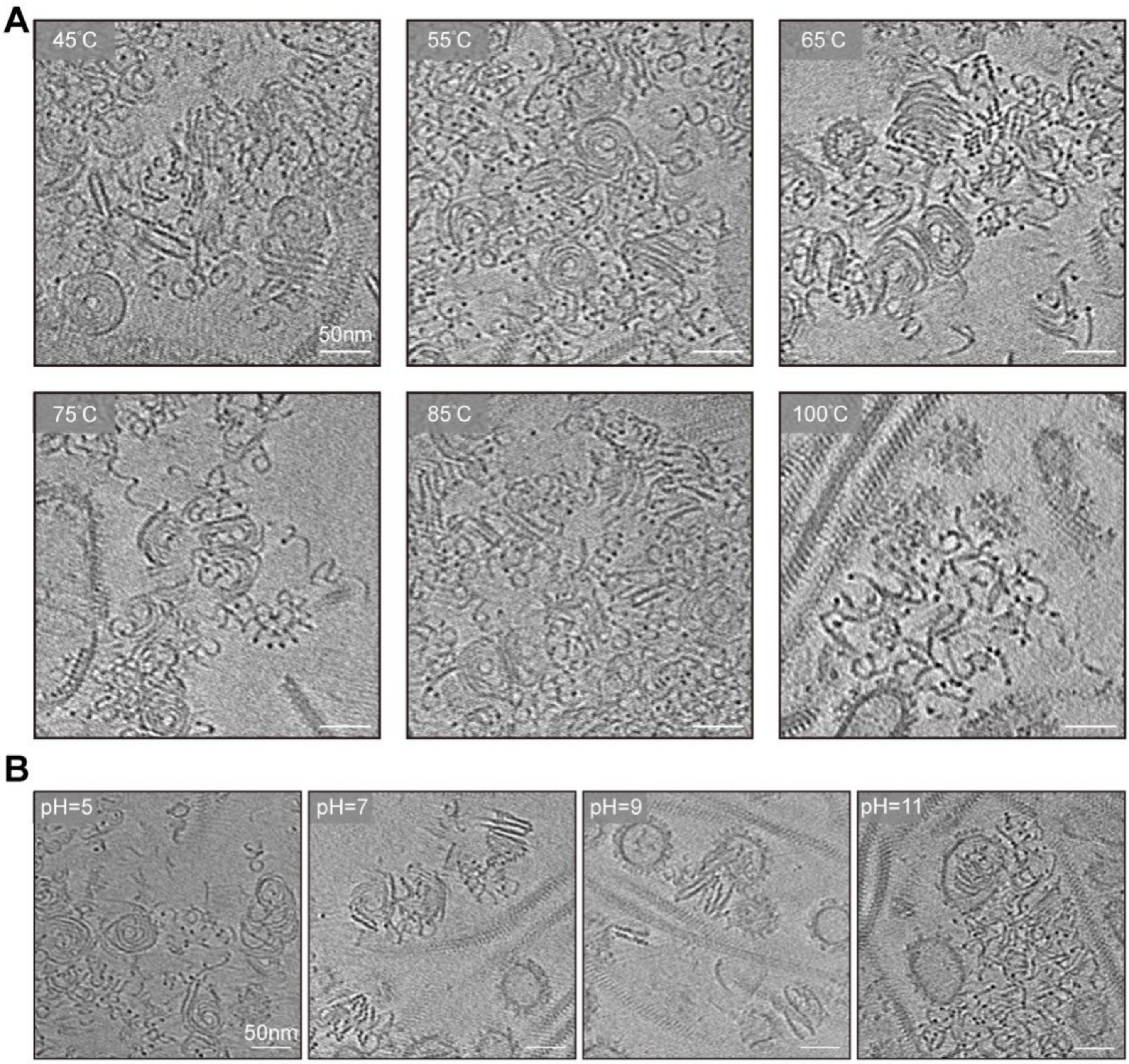


**Figure S8:** The structural changes of the naked SEV1 virions with fluctuated temperatures (A) and pH (B).

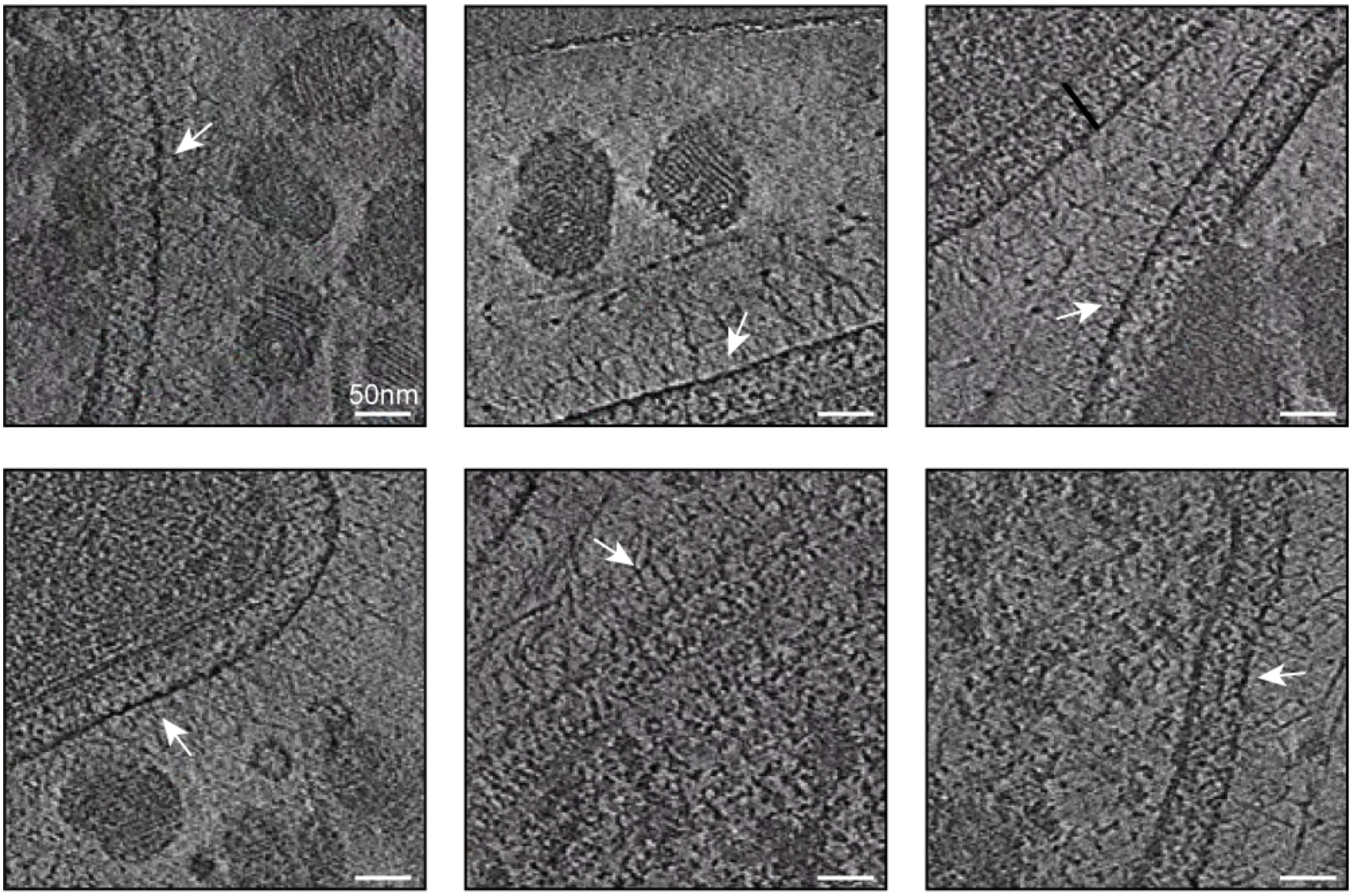


**Figure S9: Glycan chains of the S-layer.** The glycan chains are indicated by white arrows. The scale bar is 50 nm.

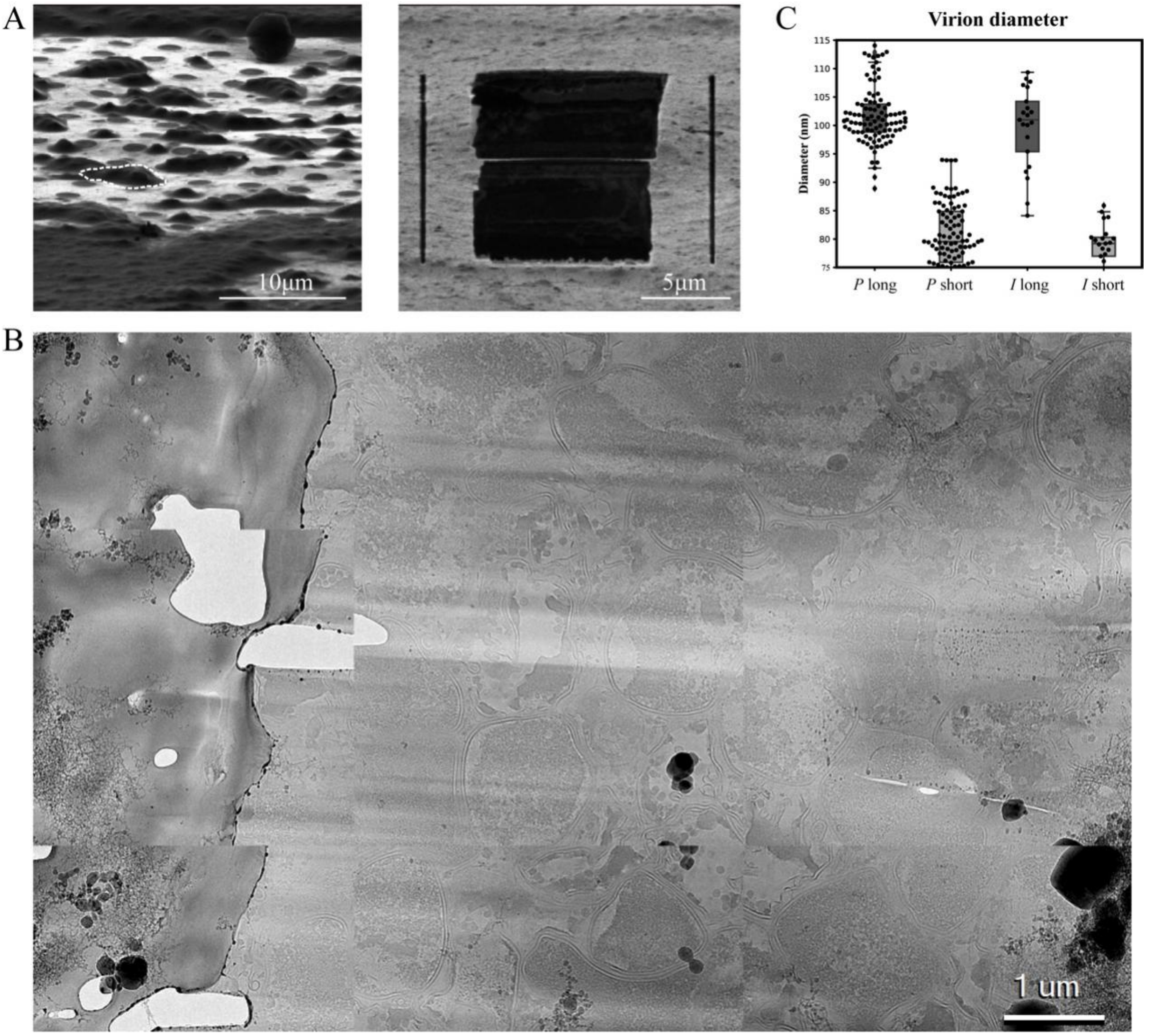


**Figure S10: The pipeline of cryo-FIB and the statistic of diameters of SEV1. (A)** SEM image of infected cells of *Sulfolobus* sp. A20 on EM grid prior to cryo-FIB milling. Example cells are outlined by dashed lines. The SEM view of the cryo-sample of the host cell. The left panel is the lamella afer cryo-FIB milling. **(B)** The projection at low magnification of infected *Sulfolobus* sp. A20 cells. **(C)** Size comparison between *in vitro* and *in situ* SEV1 virions.

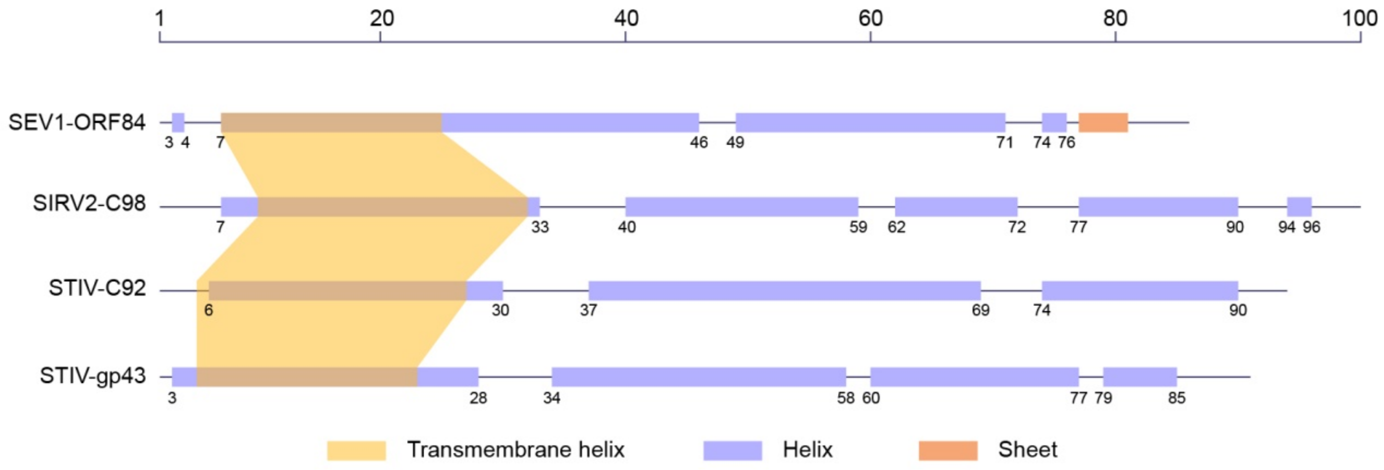


**Figure S11: Comparison of secondary structures of PVAPs.**

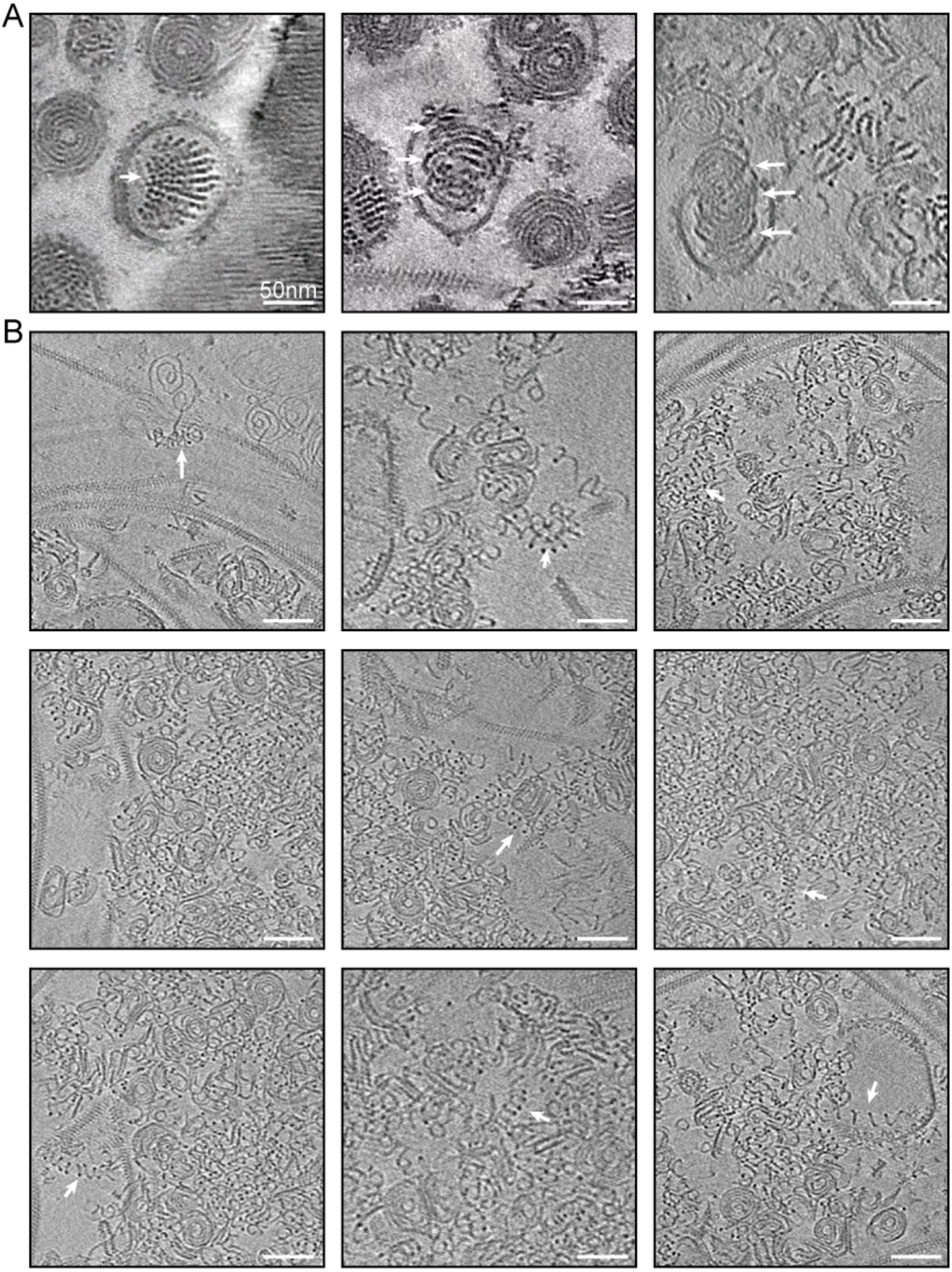


**Figure S12: Structural elements of the SEV1 nucleocapsid. (A)** The stacking discs in virions indicated by white arrows. **(B)** Single-layered helices highlighted by white arrows among the completely or partially disassembled nucleocapsids.

**Table S1. Mass spectrometry and MaxQuant generated iBAQ value**

| Protein name | Peptide sequence | iBAQ |
|---|---|---|
| SEV1-VP4 YP_009639281.1 | GVATWTAIMR;GVATWTAIMRNPEVR;INQMIANYTSVMQNMPSNADAEAR;MSRSPADVASNAWDK;NAKYTYEKTLMGVSSTGTVPITIPAQTG;NGIAGVVR;NGIAGVVRNAK;NGIAGVVRNAKYTYEK;SPADVASNAWDK;SPADVASNAWDKYLSK;TLMGVSSTGTVPITIPAQTG;YEKGVATWTAIMR;YEKGVATWTAIMRNPEVR;YTYEKTLMGVSSTGTVPITIPAQTG | 1,32E+10 |

Table S3. Functional annotation of viral proteins by NetGo.

| Name | GO | Score | Function name |
|---|---|---|---|
| vp1 | GO:0005488 | 0.918 | binding |
| vp1 | GO:0003674 | 0.918 | molecular_function |
| vp1 | GO:0005515 | 0.819 | protein binding |
| vp1 | GO:0003824 | 0.737 | catalytic activity |
| vp1 | GO:0097159 | 0.669 | organic cyclic compound binding |
| vp1 | GO:1901363 | 0.652 | heterocyclic compound binding |
| vp1 | GO:0016787 | 0.6 | hydrolase activity |
| vp2 | GO:0005488 | 0.897 | binding |
| vp2 | GO:0003674 | 0.897 | molecular_function |
| vp2 | GO:0005515 | 0.81 | protein binding |
| vp2 | GO:0003824 | 0.795 | catalytic activity |
| vp2 | GO:0016787 | 0.674 | hydrolase activity |
| vp2 | GO:0097159 | 0.669 | organic cyclic compound binding |
| vp2 | GO:1901363 | 0.623 | heterocyclic compound binding |
| vp3 | GO:0005488 | 0.904 | binding |
| vp3 | GO:0003674 | 0.904 | molecular_function |
| vp3 | GO:0005515 | 0.756 | protein binding |
| vp3 | GO:0097159 | 0.745 | organic cyclic compound binding |
| vp3 | GO:0003824 | 0.696 | catalytic activity |
| vp3 | GO:1901363 | 0.652 | heterocyclic compound binding |
| vp3 | GO:0003676 | 0.64 | nucleic acid binding |
| vp3 | GO:0016787 | 0.6 | hydrolase activity |
| vp4 | GO:0005488 | 0.918 | binding |
| vp4 | GO:0003674 | 0.918 | molecular_function |
| vp4 | GO:0005515 | 0.81 | protein binding |
| vp4 | GO:0097159 | 0.802 | organic cyclic compound binding |
| vp4 | GO:1901363 | 0.769 | heterocyclic compound binding |
| vp4 | GO:0003676 | 0.764 | nucleic acid binding |
